# Lack of an atypical PDR transporter generates an immunogenic *Cryptococcus neoformans* strain that drives a dysregulated and lethal immune response in murine lungs

**DOI:** 10.1101/2024.06.17.599354

**Authors:** Christopher J. Winski, Peter V. Stuckey, Armando M. Marrufo, Georgina Agyei, Robbi L. Ross, Tamanna Urmi, Sarah Chapman, Felipe H. Santiago-Tirado

**Affiliations:** Department of Biological Sciences, University of Notre Dame, Notre Dame, Indiana, USA; Integrated Biomedical Sciences, University of Notre Dame, Notre Dame, Indiana, USA; Eck Institute for Global Health, University of Notre Dame, Notre Dame, Indiana, USA; Warren Center for Drug Discovery, University of Notre Dame, Notre Dame, Indiana, USA; Integrated Imaging Facility, University of Notre Dame, Notre Dame, Indiana, USA

## Abstract

*Cryptococcus neoformans* is an opportunistic fungal pathogen responsible for >150,000 deaths every year with a mortality rate as high as 81%. This high medical burden is due, in part, to an incomplete understanding of its pathogenesis. In a previous study, we identified a cryptococcal atypical ATP-binding cassette (ABC) pleiotropic drug resistance (PDR) transporter, *PDR6*, that affected antifungal resistance and host interactions. Here, we follow-up on the role of *PDR6* in cryptococcal virulence. *In vivo*, mice infected with the *pdr6*Δ strain display altered symptomatology and disease progression. Specifically, we observed a significant increase in the innate immune cell populations in the *pdr6*Δ-infected mice when compared to their WT-infected littermates. Furthermore, quantification of pulmonary cytokines/chemokines revealed a robust increase of pro-inflammatory cytokines in mice infected with the *pdr6*Δ mutant strain. Despite the documented sensitivity of the *pdr6*Δ strain to azole antifungal drugs, the treatment of *pdr6*Δ-infected animals with antifungals did not affect survival, yet treatment with a corticosteroid significantly extended survival, highlighting the importance of a balanced/controlled host immune response. Results with mice that mount opposing immune responses supports out hypothesis that the *pdr6*Δ strain induces a hyper-inflammatory immune response, and that the mice succumb to immune-dependent tissue damage rather than the fungal burden. This altered immune response is driven, in part, by changes in the mutant’s surface. Taken together, this study provides insights regarding cryptococcal pathogenesis and highlights additional functions of PDR-type ABC transporters in pathogenic fungi.

**IMPORTANCE:** Yeasts of the *Cryptococcus* genus, especially *C. neoformans*, can cause disease with unacceptably high mortality. This is due to delays in diagnostics, ineffective treatments, and an incomplete understanding of the interactions between this fungus and our immune system. In this study, we expand our knowledge of the biological function of the *PDR6* gene, particularly its effect on modulating the host’s immune response. Normally, *C. neoformans*’s infections are characterized by an anti-inflammatory response that is unable to control the yeast. In the absence of *PDR6*, the response to the infection is a dysregulated pro-inflammatory response that initially controls the fungi but eventually results in death of the host due to too much tissue damage. This is due, in part, to an altered fungal surface. Given the dual role of *PDR6* in modulating antifungal sensitivity and immune responses, this work provides important insights that may lead to new or improved therapeutics.

## INTRODUCTION

Commonly associated with soil and bird guano, *Cryptococcus neoformans* is an environmental, opportunistic yeast responsible for >194,000 infections every year (1). Upon inhalation of spores or desiccated yeast, *C. neoformans* reaches the alveoli where it encounters alveolar macrophages and other host immune cells (2); however, this fungus is capable of establishing a unique intracellular niche that allows it to survive and replicate within these host cells (3). Under certain immune conditions, this neurotropic fungus can then escape the lungs, traverse the blood-brain-barrier, and enter the central nervous system (CNS), resulting in cryptococcal meningoencephalitis (CME) with a mortality rate as high as 81% (1, 4, 5). Several lines of evidence indicate that in addition to its use of host immune cells as replication havens, *C. neoformans* can use infected cells as vehicles for dissemination and brain invasion (5, 6). Hence, understanding this complex cryptococcal-phagocyte interaction from both fungal and host perspectives represents a challenging but clinically relevant avenue of research given its role in determining the course and outcome(s) of infection.

In addition to its ability to survive intracellularly, *C. neoformans* has several other virulence factors including an antiphagocytic polysaccharide capsule that provides protection from host stressors and masks several cryptococcal pathogen-associated molecular patterns (PAMPs) including chitin, chitosan, mannoproteins, and β-glucans from host phagocytes (7). The capsule, structurally composed of glucuronoxylomannan (GXM) and glucuronoxylomannogalactan (GXMGal), is constitutively secreted/shed and is known to modulate the host immune response in ways that promote infection (8). Additionally, *C. neoformans* synthesizes cell wall-localized melanin that protects the yeast from ultraviolet radiation and oxidative/nitrosative stress (9). Finally, the fungus displays thermotolerance, able to grow at physiological-relevant conditions (37°C + 5% CO_2_) (10–12). These virulence factors are integral to the pathogenesis of *C. neoformans* and as such, have been well-studied. However, *C. neoformans* utilizes additional virulence factors which remain significantly understudied or completely unknown. For example, we previously identified *PDR6* (also known as *AFR3*), a novel ATP-binding cassette (ABC) transporter of the pleiotropic drug resistance (PDR) class, and showed that it had a role in various cryptococcal virulence facets (13, 14). Present only in plants and fungi, PDR transporters hydrolyze ATP to transport substrates across biological membranes and have been primarily implicated in antifungal resistance (15, 16). Indeed, deletion of *PDR6* (*pdr6*Δ) results in hyper-susceptibility to the azole-class of antifungals, but also showed *in vitro* and *in vivo* defects in virulence. Thus, the role of *PDR6* in fungal pathogenesis and modulating the host immune response warranted further investigation.

During microbial infections, the host’s immune system is tasked with mounting an appropriate and balanced immune response that will clear the infection while ideally limiting tissue damage (17). An exaggerated Th1, or pro-inflammatory, response will likely clear a microbial infection but also result in excessive cellular and tissue damage. Opposingly, an inappropriate Th2, or anti-inflammatory, response will limit host damage but allow for uncontrolled microbial proliferation and dissemination. Many pathogens take advantage of this dichotomy to promote their infection. For example, early in HIV infection, this virus promotes the secretion of pro-inflammatory cytokines IFN-γ and TNF-α, generating a strong Th1 response that is permissive to viral replication (18, 19). In the context of *C. neoformans* infections, the fungus promotes an anti-inflammatory Th2 response that allows for intracellular survival and increased replication and dissemination, ultimately resulting in patient mortality (20, 21). It is evident, however, that between these two immune response extremes, there is a whole spectrum of responses to microbial infections, making this topic extremely complex but also an important target for therapeutics. Hence, increased understanding of the immune response against *Cryptococcus* can inform future therapeutic strategies where the immune response could be modulated to benefit the host.

Our previous work characterizing *PDR6* showed that it had important roles in cryptococcal biology and suggested that it was necessary for full virulence (13). Here, we study the role of *PDR6* in the establishment and progression of disease using a murine inhalation model of cryptococcosis. We show that a lower inoculum or treatment with antifungal drugs, significantly decreased *pdr6*Δ fungal burdens, but do not change the outcome of the infection by this mutant. Instead, the survival studies and immunological analyses suggest that the *pdr6*Δ mutant elicits an altered host immune response characterized by significant increases in pro-inflammatory cytokines, chemokines and innate immune cells. We treated the *pdr6*Δ-infected mice with a corticosteroid and significantly extended their survival despite little to no change in fungal burden. Moreover, using mouse breeds that are known to establish opposing immune responses, we show that indeed the host’s immune response is the main determinant of the *pdr6*Δ infection outcome. This dysregulated immune response might be a consequence of fungal surface alterations that expose antigenic and immunogenic components that normally would be hidden from the immune system. Collectively, this study furthers our understanding of cryptococcal pathogenesis, highlights the clinical importance of the host immune response in controlling fungal infections, while also providing novel functions and insights of fungal PDR-type transporters in the context of disease progression.

## RESULTS

### *pdr6*Δ-infected mice succumb to infection despite decreased inoculum and fungal burden

In our previous study we determined that *PDR6* was required for complete cryptococcal virulence as mice infected with the *pdr6*Δ mutant lived significantly longer and had decreased fungal burdens when compared to their H99-infected littermates (13). Consistently, this finding was recapitulated by *Oliveira et al.* using the *Galleria mellonella* invertebrate infection model (14). Notably, the *pdr6*Δ-infected animals displayed altered symptomatology compared to their H99-infected littermates. Specifically, at endpoint, the WT-infected mice displayed altered behavior associated with neurological deficiencies, traditional of cryptococcal meningitis; in contrast, the *pdr6*Δ-infected mice exhibited respiratory symptoms characterized by rapid and labored breathing, suggestive of pneumonia. We hypothesized that the altered symptomatology could be explained, in part, by differences in the establishment and progression of disease. Previously, fungal burdens were only quantified at time of death (TOD), potentially overlooking differences earlier on in the infection. To obtain a comprehensive view of the infection over time, we inoculated mice with 5 x 10^4^ cells of H99 or *pdr6*Δ strains, and quantified fungal burden every seven days over the course of the infection (Fig. 1A). Interestingly, we found that at 7 days post infection (dpi) there were no differences in the fungal burdens between the H99 and *pdr6*Δ-infected animals in all organs tested, suggesting that the strains established infection similarly. Following this initial establishment, however, at 14 dpi there was a significant decrease in fungal burden for the *pdr6*Δ strain compared to the WT. At 17 dpi, the H99-infected mice succumbed to the infection with average lung, brain, and spleen burdens of 3.7 x 10^8^, 1.9 x 10^6^, and 1.3 x 10^4^ CFU while the *pdr6*Δ-infected mice CFUs were 3.6 x 10^7^, 5.1 x 10^3^, and 677, respectively. Between 14 and 28 dpi, the fungal burden of the *pdr6*Δ-infected animals remains relatively constant in the brain and spleen, with a gradual, although small, increase in the lungs. Then, after 28 dpi, there are gradual increases in all organs with final average CFU counts of 5.5 x 10^7^, 1.1 x 10^5^, and 1.5 x 10^3^ in the lungs, brain, and spleen, respectively. The differences in CFUs highlight key differences between H99 and *pdr6*Δ disease progression and suggests a potential role of the adaptive immune response given the changes at 14 dpi.

**Fig. 1.**
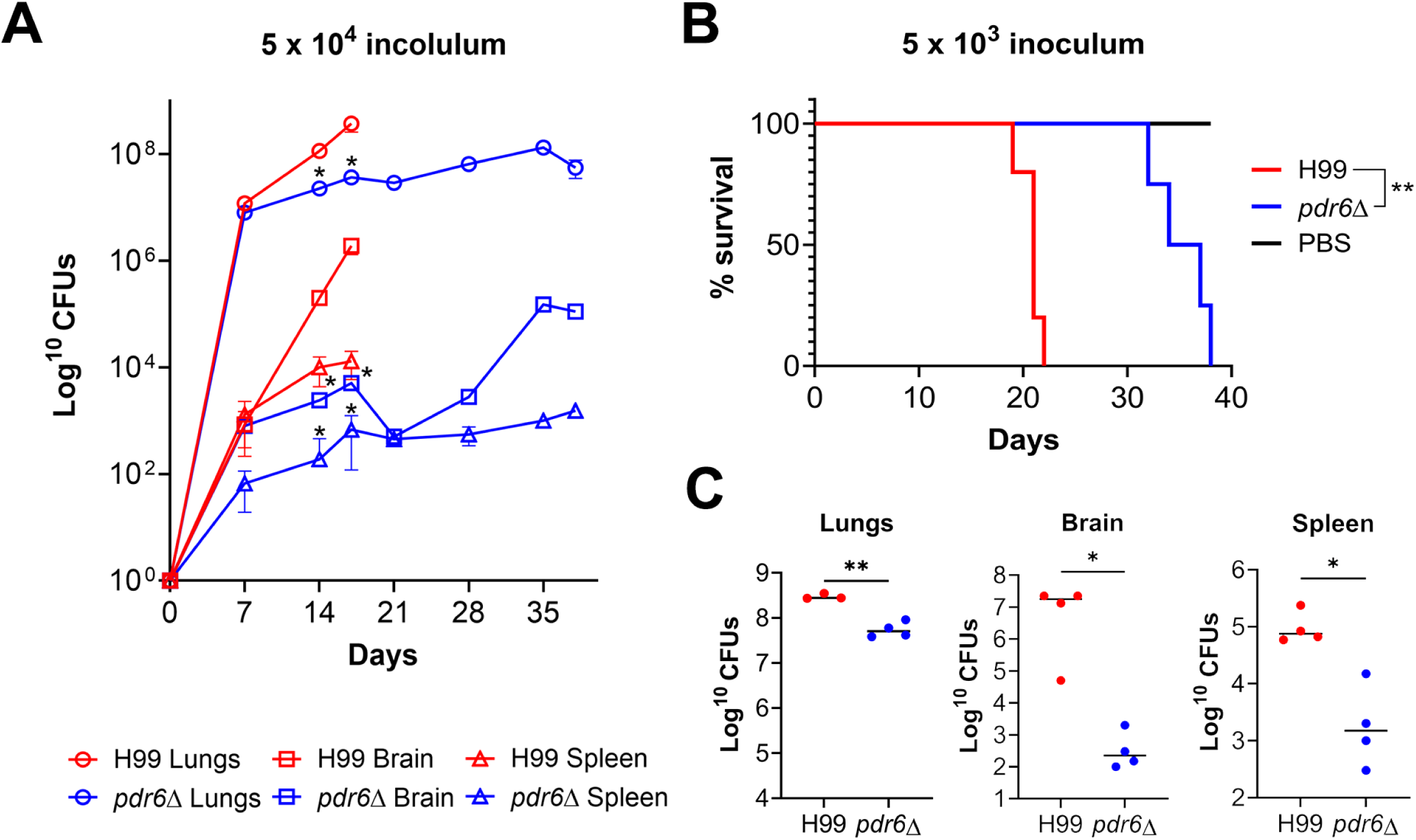
*C. neoformans pdr6*Δ-infected mice succumb to infection despite decreased fungal burden. **(A)** Quantification of fungal burden over time to monitor disease progression. Mice were inoculated with 5 x 10^4^ cells and sacrificed every 7 days and organs were harvested to determine CFUs. Significance was determined using Mann-Whitney test, *P<0.05. **(B)** Survival studies of animals infected with H99 and *pdr6*Δ strains, or mock-infected (PBS). 4 A/J mice per group were intranasally infected with 5 x 10^3^ cryptococcal cells or mock infected with sterile PBS and monitored for 38 days (experiment was terminated when the last infected mouse reached endpoint). Significance was determined using Mantel-Cox test, **P<0.01. **(C)** Quantified organ burdens of H99 and *pdr6*Δ-infected mice at TOD. Organs were harvested, homogenized, and dilutions were plated on YPD. Plates were incubated for 48 h and CFUs were counted. Each data point represents a mouse and black bars represent the mean. Significance was determined using Mann-Whitney test, *P<0.05; **P<0.01.

Experimentally, the mice in our previous studies were inoculated with 5 x 10^4^ cells, a common inoculum for the murine inhalation model of cryptococcosis. Given the *in vitro* defects of the *pdr6*Δ strain (13), and the fact that initially, the mice were able to control the infection (Fig. 1A), we hypothesized that by lowering the fungal inoculum, the survival of the *pdr6*Δ-infected mice would be extended with a potential for complete clearance of the infection. To assess this, we decreased the initial inoculum 10-fold and infected mice with 5 x 10^3^ cells and monitored survival (Fig. 1B). The H99-infected mice had a mean survival of 21 days and the *pdr6*Δ-infected mice displayed a mean survival of 36 days, similar to our studies with 10-fold higher inoculum. Quantification of the fungal burden showed similar CFUs in all organs between the animals infected with low and high WT inoculum (Fig. 1C compared to 1A); however, despite having similar lungs CFUs, the animals infected with low *pdr6*Δ inoculum had lower brain and spleen CFUs at TOD relative to the higher inoculum (Fig. 1C compared to 1A). Evidently, using a lower inoculum did not change the trend we have seen before: the *pdr6*Δ-infected mice still survive longer with significant decreases in fungal burden across all organs. However, notably, in the *pdr6*Δ-infected mice the lower inoculum results in lower overall organ burdens, with less dissemination, yet the *pdr6*Δ-infected mice still succumbed to the infection with the same dynamics as when using a higher inoculum (35-38 days; Fig. 1B).

### Infection with *pdr6*Δ results in increased tissue damage

Despite initially colonizing the pulmonary environment, the *Cryptococcus*-associated fatal pathology occurs in the brain due to the meningoencephalitis (22, 23). The altered symptomatology coupled with the differential disease progression suggests that the *pdr6*Δ strain likely affects the host immune response. Furthermore, when performing the previous fungal burden experiments, we observed significant phenotypic differences in lung sizes in H99 v. *pdr6*Δ-infected mice (Fig. 2). To qualitatively assess any potential differences at the cellular and tissue level, we harvested the lungs of mock, H99, and *pdr6*Δ-infected mice, stained with hematoxylin and eosin (H&E), and assessed tissue damage (Fig. 2 and Table 1). At 14 dpi, the gross anatomical images clearly show increased size and inflammation of the H99-infected lungs when compared to the *pdr6*Δ-infected. Additionally, the 10X and 40X images display deterioration of the lung parenchyma and infiltration of *C. neoformans* which is not observed in the *pdr6*Δ-infected lungs (Fig. 2A). At 35 dpi, the *pdr6*Δ-infected lungs are severely inflamed, infiltrated with yeast, and characterized by the presence of many inflammatory immune cells in the peri-bronchial and perivascular space (Fig. 2B). Table 1 provides a quantitative summary comparing the H99 and *pdr6*Δ-infected pulmonary tissue over time. Consistent with our observations in the fungal burden experiments, there were little to no differences at 7 dpi. However, at 14 dpi the *pdr6*Δ-infected lung contains the yeast in granuloma-like regions and have 30% less *C. neoformans* than the WT-infected tissues. Furthermore, an increase in lymphocytes is observed in the *pdr6*Δ-infected lungs relative to WT (105 to 26, respectively; Table 1). Finally, as the *pdr6*Δ-infected mice approach TOD, the fungal infiltration, inflammation, and number of inflammatory immune cells in the lung tissue all increase to levels well above those observed in the WT-infected animals at TOD. Altogether, these findings suggest that the *pdr6*Δ strain elicits a differential immune response in the pulmonary environment thus affecting disease progression and symptomatology.

**Fig. 2.**
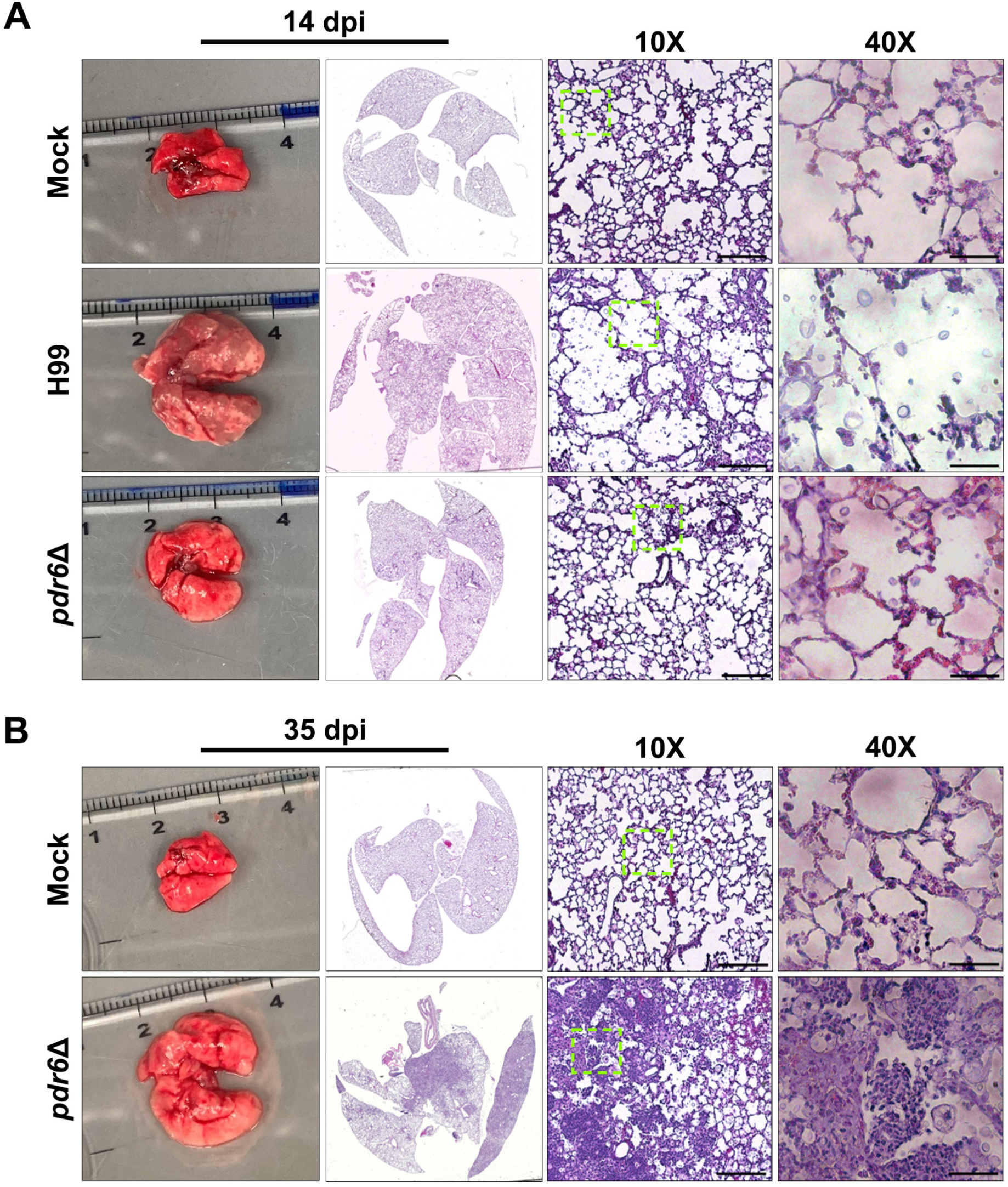
Infection with *C. neoformans pdr6*Δ results in altered histopathology. **(A &B)** Dissected lungs of female A/J mice infected with 5 x 10^4^ cells of the WT strain (H99) or *pdr6*Δ mutant strain, or mock-infected, were harvested at predetermined endpoints of 14 **(A)** and 35 **(B)** dpi. Gross organ examination revealed significant changes in inflammation and tissue damage. H&E staining was used to visualize microscopic lung pathology (inset depicted by the green boxes is shown in the 40X column). The 10X scale bar is 200 μm and the 40X scale bar is 50 μm.

**Table 1.** Quantitative analysis of H99 and *pdr6*Δ-infected lung sections over time. *All H99-infected mice had succumbed to infection by this timepoint. Histiocytes refer to leukocytes of the monocyte/macrophage/dendritic cell lineage. hpf, high power field.

| DPI | Strain | % inflammation | Fungal infiltration | Quantitation of inflammatory immune cells/hpf 400X | Observations |
| --- | --- | --- | --- | --- | --- |
| 7 | H99 | 5 – 10% | 20% alveoli contain encapsulated yeast | 8 neutrophils<br>33 lymphocytes<br>40 histiocytes | Immune cells in peri-bronchial and perivascular space. |
|  | *pdr6Δ* | 5 – 10% | 15 – 20% alveoli contain encapsulated yeast | 9 neutrophils<br>26 lymphocytes<br>19 histiocytes | Immune cells in peri-bronchial and perivascular space. Some yeast has been phagocytized by histiocytes. |
| 14 | H99 | 10% | 80 – 90% alveoli contain encapsulated yeast | 52 neutrophils<br>26 lymphocytes<br>57 histiocytes | Immune cells in peri-bronchial and perivascular space. |
|  | *pdr6Δ* | 10% | 50 – 60% alveoli contain encapsulated yeast | 43 neutrophils<br>105 lymphocytes<br>71 histiocytes | Immune cells in peri-bronchial and perivascular space. Many histiocytes have phagocytized yeast. Areas of accumulated yeast/immune cells but no true parenchymal granulomas. |
| 21* | *pdr6Δ* | 10% | 90% alveoli contain encapsulated yeast | 50 neutrophils<br>49 lymphocytes<br>55 histiocytes | Immune cells in peri-bronchial and perivascular space. |
| 28* | *pdr6Δ* | 5 – 10% | 80% alveoli contain encapsulated yeast | 8 neutrophils<br>33 lymphocytes<br>55 histiocytes | Immune cells in peri-bronchial and perivascular space and single parenchymal granuloma (1.2mm) |
| 35* | *pdr6Δ* | 60 – 70% | 100% present throughout | Too many to quantify | Consolidated parenchyma with many coalescent granulomas (histiocytes w/phagocytized yeast) and nests of neutrophils and lymphocytes with necrosis. |

### Fluconazole treatment improves fungal burden but not outcome in *pdr6*Δ-infected mice

Currently, there a very few antifungal drugs available to treat cryptococcal infections and the compounds that are available are either not effective, extremely expensive, or associated with deleterious side effects (24). Fluconazole (FLC), a fungistatic azole class antifungal, is the mainstay treatment option in low-resource regions where cryptococcal infections are endemic. Mechanistically, FLC inhibits lanosterol 14α-demethylase, an enzyme in the ergosterol biosynthesis pathway encoded by *ERG11* (25). In our previous study, we showed that the deletion of *PDR6* increased the susceptibility of *C. neoformans* to azole class antifungals including FLC (13). Given this phenotype, we hypothesized that treatment of *pdr6*Δ*-*infected mice with FLC would result in total clearance of the infection, preventing the lethal immune pathology. To test this, mice infected with either the WT or *pdr6*Δ strains, or mock infected, were treated daily (starting 1 dpi) with either a low (8 mg/kg) or high dose (16 mg/kg) of FLC and monitored for survival (Fig. 3A). The FLC treatment significantly extended survival in both the H99 and *pdr6*Δ-infected animals compared to the untreated controls, albeit to the same extent. Numerically, the mean survival for the untreated H99-infected mice was 17 days and the low and high doses extended survival 4.5 and 6 days, respectively. Similarly, the mean survival for the untreated *pdr6*Δ-infected mice was 34 days and the low and high doses extended survival 5 and 7 days, respectively. Surprisingly, although the survival was similarly extended between strains, quantification of the fungal burden revealed significant differences (Fig. 3B-G). At 14 dpi, both doses of the FLC treatment significantly decreased the fungal burden in the lungs, brain, and spleen only in the *pdr6*Δ-infected mice (Fig. 3B-D). Similarly, at TOD, the daily FLC treatment resulted in an additional substantial decrease in the fungal burden of the *pdr6*Δ-infected mice across all three organs, with several mice where the brain and spleen CFU counts were below the limit of detection (LOD) (Fig. 3E-G). Not surprisingly, neither of the FLC doses had a significant effect on WT-infected mice. This observation is interesting because despite significantly decreasing the fungal burden with the FLC treatment, the *pdr6*Δ-infected animals still succumbed to the infection.

**Fig. 3.**
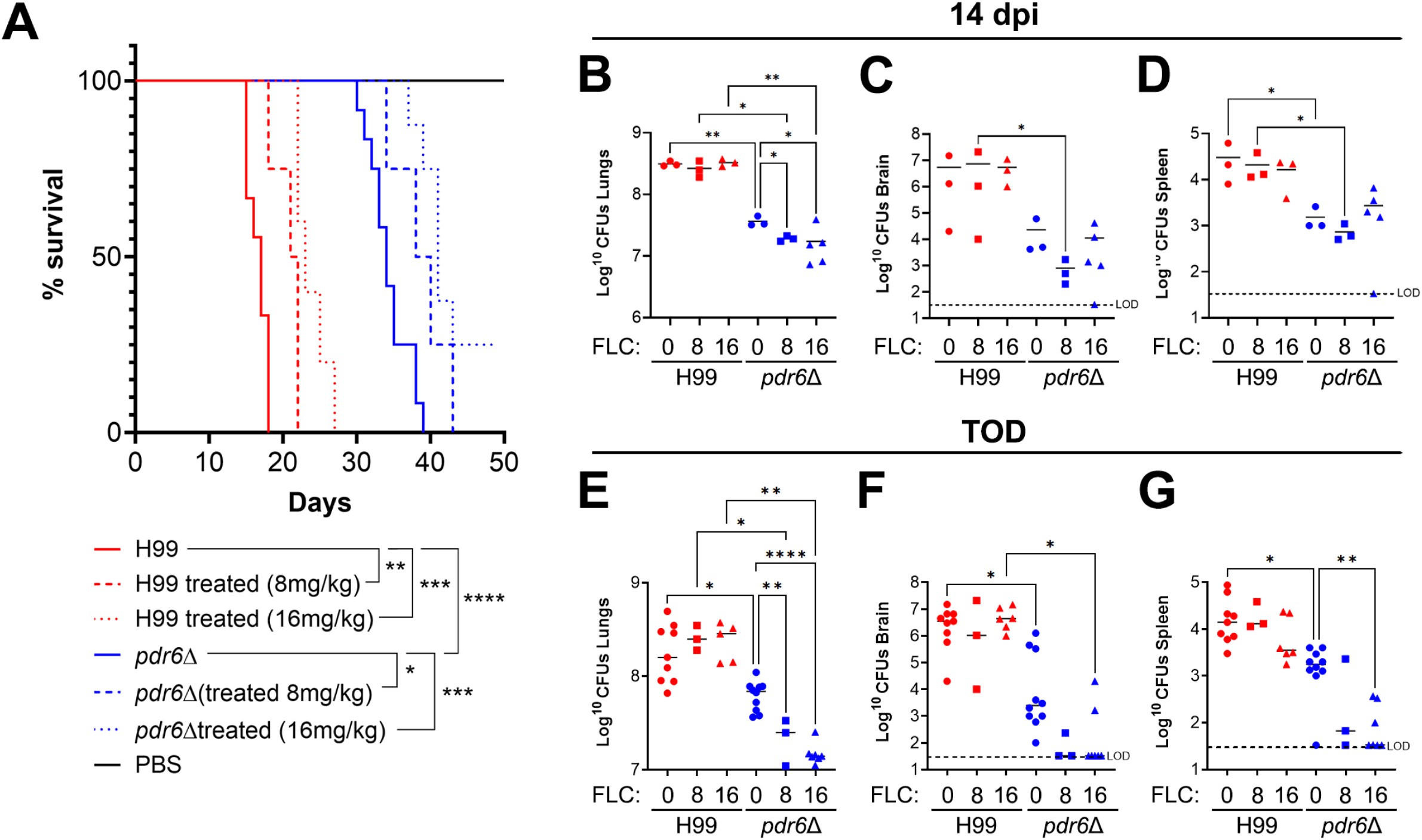
FLC treatment significantly reduces CFUs in *pdr6*Δ-infected mice without changing the outcome. **(A)** Survival curve of H99 and *pdr6*Δ-infected mice treated daily with either 8 or 16 mg/kg FLC. At least 4 A/J mice per group were intranasally infected with 5 x 10^4^ cells, or mock-infected with DPBS and monitored for up to 50 dpi (mock-infected and FLC-treated mice not shown for clarity). Significance was determined using Mantel-Cox test, *P<0.05; **P<0.01; ***P<0.001; ****P<0.0001. **(B-G)** Calculated organ burdens at 14 dpi **(B-D)** or endpoint **(E-G)**. Organs were harvested, homogenized, and dilutions were plated on YPD. Plates were incubated for 48 hr and CFUs were enumerated. Each data point represents a mouse and black bars represent the mean. Significance was determined using one-way ANOVA with multiple comparisons (Welch’s corrected), *P<0.05; **P<0.01; ****P<0.0001.

To understand how the FLC treatment was affecting the pulmonary environment, we prepared slides and performed qualitative histological analysis at TOD for both WT and *pdr6*Δ-infected mice (Fig. 4 and Table 2). The H99-infected lungs at 14 dpi treated with FLC displayed similar features to the previously pictured H99 untreated lungs, namely a high percentage of fungal infiltration and tissue breakdown. In contrast, there was a clear decrease in fungi in the FLC-treated *pdr6*Δ-infected lungs at this timepoint. Consistently, the FLC-treated *pdr6*Δ-infected lungs at 35 dpi appear to have the infection controlled in granuloma-like regions (Fig. 4, inset 2) with widespread immune cell infiltration (Fig. 4, inset 1). Interestingly, despite reduced pulmonary CFUs (Fig. 3 and Table 2) and decreased inflammation (Fig. 4 and Table 2), the *pdr6*Δ-infected mice were still succumbing to the infection. Together, these data suggest that the mice infected with the *pdr6*Δ-strain are not necessarily dying directly because of the *C. neoformans* infection, but rather indirectly because of an inappropriate host immune response.

**Fig. 4.**
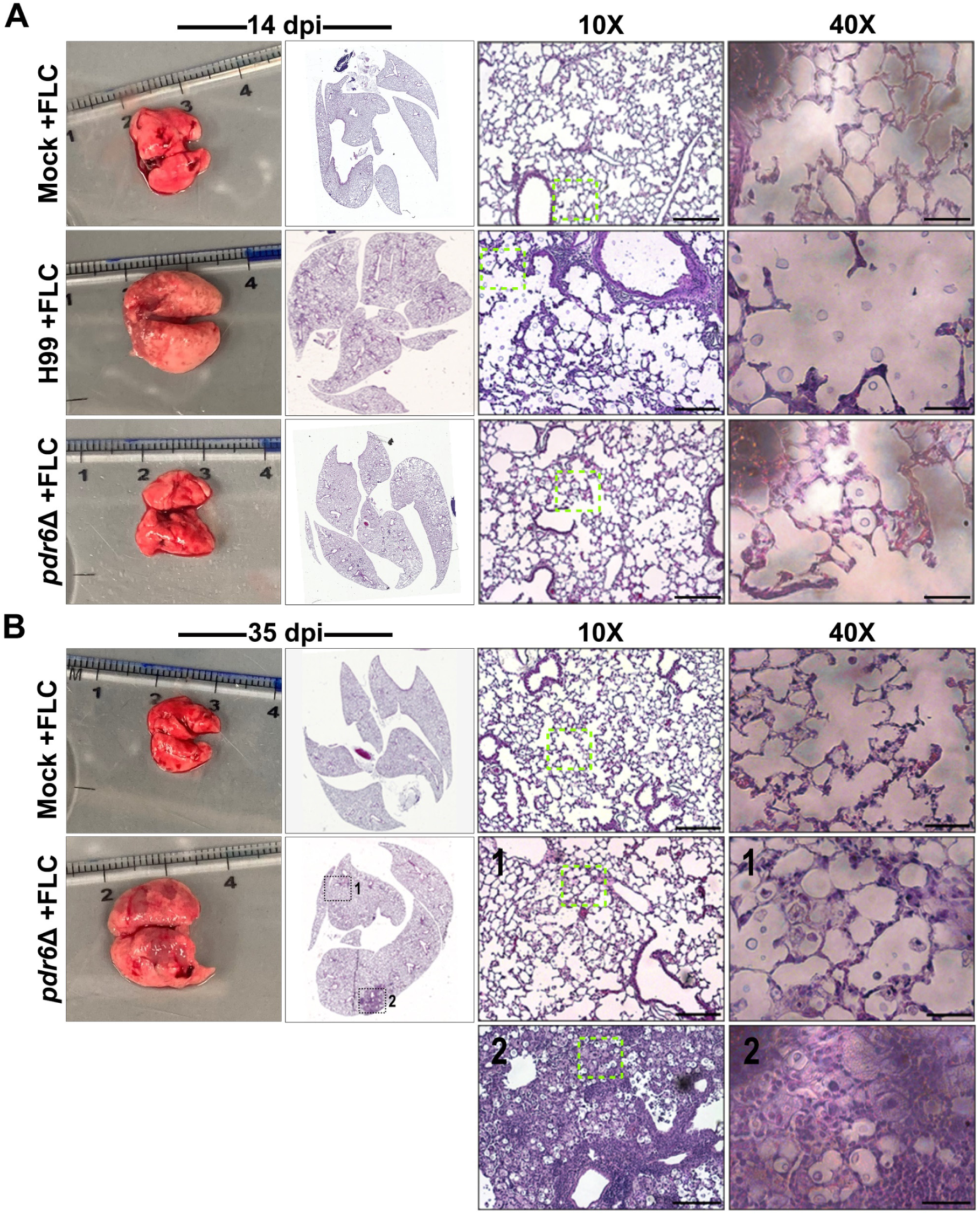
FLC treatment improves lung pathology in *pdr6*Δ-infected animals. Dissected lungs of female A/J mice infected with 5 x 10^4^ cells of the WT strain (H99) or *pdr6*Δ mutant strain and treated daily with 16 mg/kg FLC were harvested at 14 and 35 dpi. Gross organ examination revealed alterations in fungal containment and disease progression. H&E staining was used to visualize microscopic lung pathology. Insets 1 & 2 highlight differential histopathology in the FLC-treated *pdr6*Δ-infected mice at 35 dpi (inset depicted by the green boxes is shown in the 40X column). Inset 1 shows an intact lung parenchyma with several fungi and immune cells while inset 2 is characterized as a granuloma. The 10X scale bar is 200 μm and the 40X scale bar is 50 μm.

**Table 2.** Quantitative analysis of Fluconazole treated H99 and *pdr6*Δ-infected lung sections over time. Histiocytes refer to leukocytes of the monocyte/macrophage/dendritic cell lineage. hpf, high power field.

| DPI | Strain | % inflammation | Fungal infiltration | Quantitation of inflammatory immune cells/hpf 400X | Observations |
| --- | --- | --- | --- | --- | --- |
| 14 | Mock +FLC | 0% | None | - | - |
|  | H99 +FLC | 5 – 10% | ~ 60% alveoli contain encapsulated yeast | 50 – 60 neutrophils<br>40 – 50 lymphocytes<br>15 histiocytes | Moderate number of inflammatory cells predominately present in perivascular & peribronchial areas. |
|  | *pdr6Δ* +FLC | < 5% | 30 – 40% alveoli contain encapsulated yeast | 1 – 5 neutrophils<br>0 – 15 lymphocytes<br>1 – 5 histiocytes | Mild infiltrate of inflammatory cells in focal peribronchial & perivascular areas. |
| 35 | Mock +FLC | 0% | None | - | - |
|  | *pdr6Δ* +FLC | 5 – 10% | 45 – 50% alveoli contain encapsulated yeast | 5 – 10 neutrophils<br>10 – 20 lymphocytes<br>< 5 histiocytes | Mild inflammatory cell infiltrate in many peribronchial and perivascular areas.<br>Consolidation of adjacent alveoli in solid areas. |

### *C. neoformans pdr6*Δ strain elicits a strong pro-inflammatory immune response

During microbial infections, the host’s immune response ultimately determines the clinical outcome (26). Ideally, a balanced Th1 pro-inflammatory and Th2 anti-inflammatory immune response will allow for microbial clearance and subsequent tissue repair. Previous studies have determined that *C. neoformans* skews the immune response towards a permissive anti-inflammatory Th2 response, thus benefiting the yeast and allowing for fungal replication and subsequent dissemination (27). Given our previous observations, we hypothesize that the *pdr6*Δ strain is instead generating a strong pro-inflammatory immune response that is able to control the infection initially, but also damaging large areas of pulmonary tissue. To assess this, we quantified the secreted cytokines and chemokines profile in the lungs and brains of H99 and *pdr6*Δ-infected mice at their respective TODs (Fig. 5 and Fig. S1). Interestingly, we found that several of the pro-inflammatory cytokines are significantly upregulated in the *pdr6*Δ-infected lungs relative to WT (Fig. 5A). Specifically, the production of IL-1α, IL-1β, and MIP-1β (CCL4) is increased 13-fold, 25-fold, and 37-fold respectively, when compared to the lungs of the H99-infected mice. Interestingly, the expression of IL-13 and IL-33, anti-inflammatory cytokines associated with allergic diseases, where also statistically upregulated in the *pdr6*Δ-infected mice relative to WT. A similar trend was observed in the brain, although relative to the pulmonary tissues, it was more muted (Fig. 5B), with statistically significant upregulation of G-CSF and IL-3 relative to the H99-infected brain. This weakened expression of inflammatory cytokines in the brain might be due to the decreased capacity of the *pdr6*Δ mutant to disseminate coupled with the immune-privileged status of the CNS. During a typical cryptococcal infection, the expression of anti-inflammatory cytokines and chemokines are increased thus generating a Th2 response, however, in the context of an infection with the *pdr6*Δ strain, the secretion of pro-inflammatory cytokines and chemokines is significantly increased resulting in a damaging Th1 immune response.

**Fig. 5.**
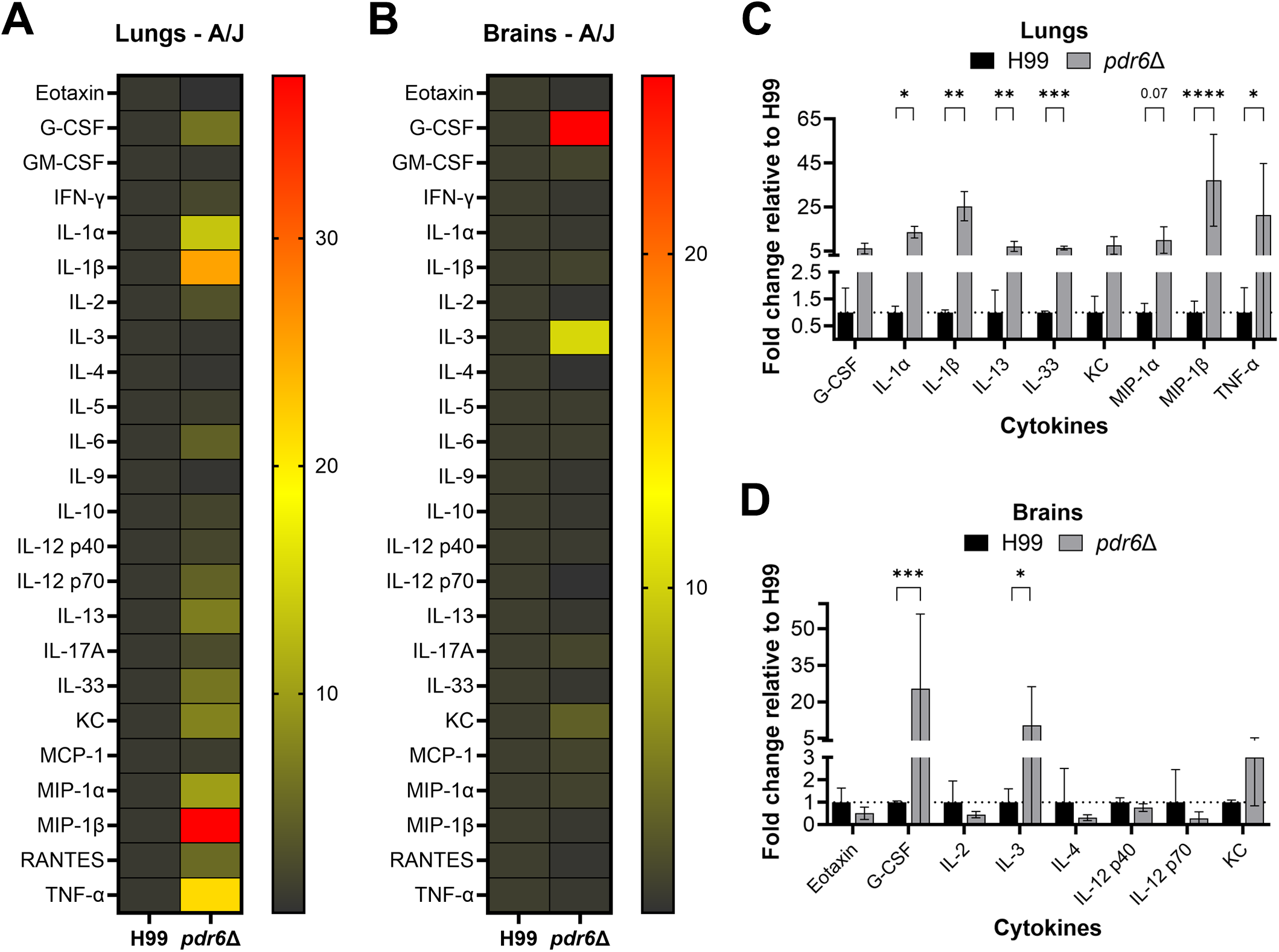
Infection with the *pdr6*Δ mutant elicits a robust pro-inflammatory immune response. **(A &B)** Heatmap of the pro-inflammatory and anti-inflammatory cytokines/chemokines in the lungs **(A)** and brains **(B)** of *C. neoformans*-infected mice. Female A/J mice were intranasally infected with 5 x 10^4^ cells of the H99 or *pdr6*Δ strain and euthanized at endpoint. Homogenates from the lungs and brain of at least 4 mice per group were used to perform Proteome Profiler Mouse XL Cytokine Arrays and Bio-Plex Multi-Plex Arrays. **(C & D)** Expression of selected cytokines/chemokines in the lungs **(C)** and brains **(D)** of *pdr6*Δ-infected mice was normalized to H99 (dotted line). Significance was determined by Ordinary 2-way ANOVA with multiple comparisons, *P<0.05; **P<0.01; ***P<0.001; ****P<0.0001.

### Corticosteroid treatment extends survival in *pdr6*Δ-infected mice

Clinically, corticosteroids like dexamethasone are prescribed to downregulate the immune system. Mechanistically, these anti-inflammatory drugs function by suppressing the migration of neutrophils and decreasing lymphocyte colony proliferation (28). The potential of dexamethasone as a treatment strategy during cryptococcal infection was investigated in a clinical trial but was prematurely concluded because of adverse effects and increased mortality (29, 30); however, corticosteroids have been used successfully to treat CME-associated inflammatory disorders such as postinfectious inflammatory response syndrome and immune reconstitution inflammatory syndrome (IRIS)(22). Given the robust pro-inflammatory immune response observed in the *pdr6*Δ-infected mice, akin to fatal IRIS in humans, we hypothesized dexamethasone treatment would dampen the aggressive Th1 response resulting in fungal clearance and improved survival. Experimentally, we infected mice with either the H99 or *pdr6*Δ strain, or mock infected, and treated them daily (starting at 1 dpi) with either 0.3 mg/kg dexamethasone or a dexamethasone and 16 mg/kg FLC combination and assessed survival (Fig. 6A). The H99-infected mice that received the dexamethasone treatment displayed a significant reduction in survival (mean survival of 13 days vs 17 days) when compared to the untreated H99 control, while the combo therapy restored survival to that of the untreated H99-infected group (mean survival of 16 days vs 17 days). This result corroborates the corticosteroid finding reported in the human clinical trial (29, 30). In terms of the *pdr6*Δ-infected mice, both the dexamethasone alone and in combination with FLC significantly extended survival. Numerically, the mean survival of the untreated *pdr6*Δ-infected control group was 34 days and was extended to 55 and 58 days in the dexamethasone and combination treated groups, respectively. The survival studies were stopped after 60 dpi because the remaining mice, including the mock-infected receiving the daily dexamethasone treatment (not shown), appeared sick despite not meeting endpoint criteria. We concluded that after two months of daily corticosteroid injections the immune systems of the mice had been ablated resulting in their poor appearances. Despite the dexamethasone treatment extending survival in the *pdr6*Δ-infected animals, the fungal burdens were differentially affected over the course of the experiment (Fig. 6B-G). Specifically, at 14 dpi the addition of dexamethasone significantly decreased the fungal burden in the lungs in both the H99 and *pdr6*Δ-infected mice and little to no changes were observed in either strain in the brain or spleen CFUs (Fig. 6B-D). At TOD the dexamethasone treatment in the *pdr6*Δ-infected animals significantly increased the lung CFUs and the combination therapy returned it to levels comparable to the untreated *pdr6*Δ-infected control (Fig. 6E). Although not statistically significant, a similar trend was observed in the brain and spleen of the *pdr6*Δ-infected mice (Fig. 6F & G). These effects were not seen in the WT-infected mice. The dynamic changes in fungal burdens in the lungs of the dexamethasone and combination treated *pdr6*Δ-infected mice can be explained by the absence or lack of inflammatory immune cells to control the infection and the addition of FLC lowers the CFUs as the fungistatic nature of the antifungal is unable to clear the initial infection. Collectively, these data further confirm and highlight the altered disease progression elicited by the *pdr6*Δ mutant and, more importantly, demonstrate the positive clinical outcome associated with controlling the inappropriate pro-inflammatory immune response.

**Fig. 6.**
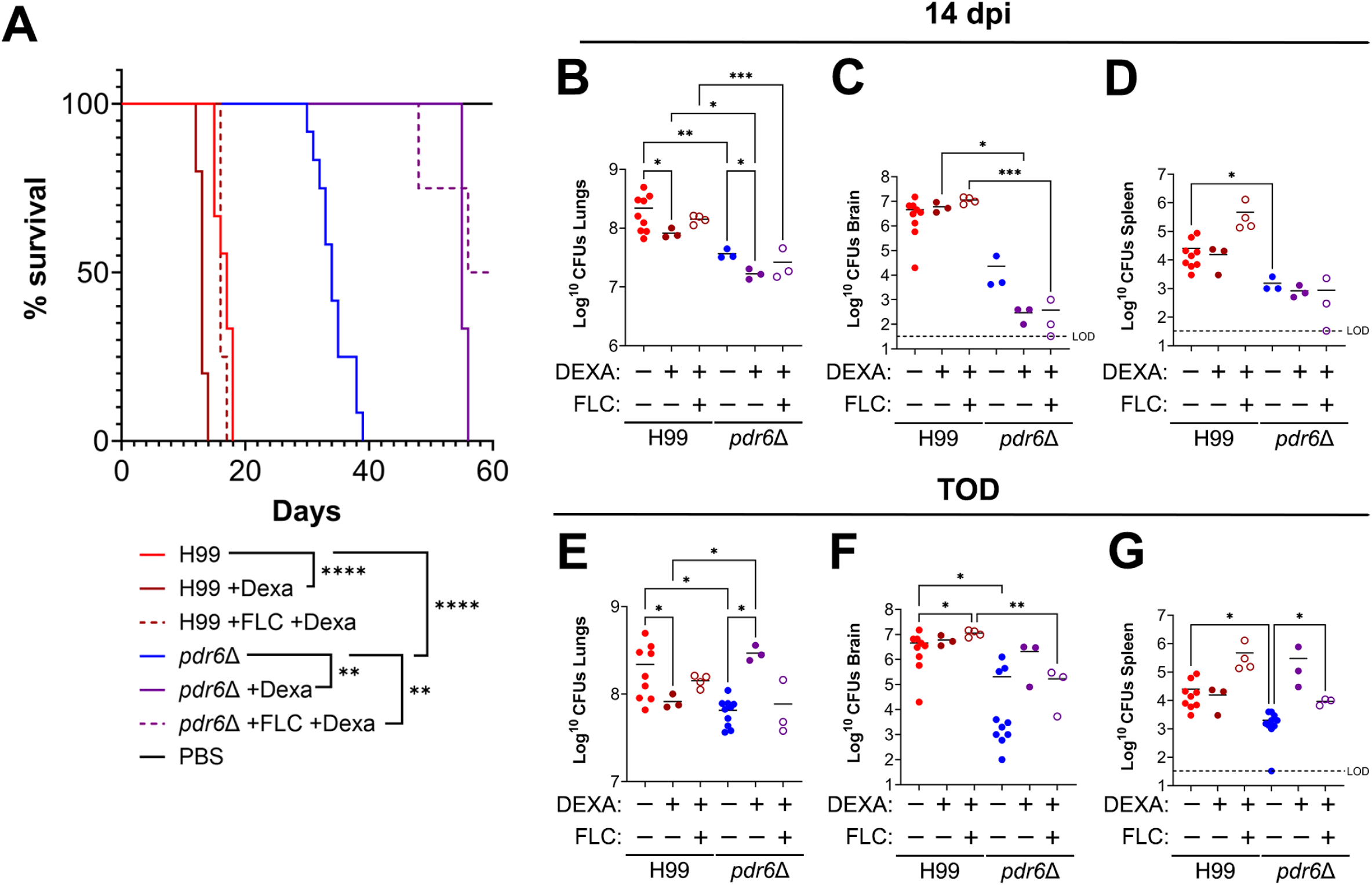
Dexamethasone treatment significantly extends survival of *pdr6*Δ-infected mice. **(A)** Survival curve of H99 and *pdr6*Δ-infected mice treated daily with either 0.03 mg/kg Dexa or 0.03 mg/kg Dexa and 16 mg/kg FLC. At least 3 A/J mice per group were intranasally infected with 5 x 10^4^ cells, or mock-infected with DPBS and monitored for up to 60 dpi (mock-infected and treated mice not shown for clarity). Significance was determined using Mantel-Cox test, **P<0.01; ****P<0.0001. **(B-G)** Calculated organ burdens at 14 dpi **(B-D)** or endpoint **(E-G)**. Organs were harvested, homogenized, and dilutions were plated on YPD. Plates were incubated for 48 hr and CFUs were enumerated. Each data point represents a mouse and black bars represent the mean. Significance was determined using one-way ANOVA with multiple comparisons (Welch’s corrected) *P<0.05; **P<0.01; ***P<0.001.

### Combination dexamethasone and fluconazole treatment reduces pulmonary tissue damage in *pdr6*Δ-infected animals

Based on our previous results, it is evident that the addition of the corticosteroid is benefiting the *pdr6*Δ-infected mice by significantly extending survival. Histological analysis of the lung tissue at TOD for both WT and *pdr6*Δ-infected mice was performed to visualize changes in fungal infiltration, tissue inflammation, and immune cell recruitment (Fig. 7 and Table 3). Visually, the treatment with dexamethasone had a profound effect given the qualitative reduction in inflammatory cells across all conditions. The lungs of the H99-infected mice that received dexamethasone are not as inflamed as those from untreated mice but the destruction of the lung tissue is devastating (Fig. 7A). The lack of immune cells allowed for the uncontrolled proliferation of *C. neoformans* resulting in increased damage of the pulmonary tissue. Similarly, the dexamethasone treated lungs of the *pdr6*Δ-infected display intense fungal proliferation but there is qualitatively less damage when compared to the H99-infected animals (Fig. 7B and Table 3). The combination dexamethasone and FLC treatment reduced the fungal infiltration in both the H99 and *pdr6*Δ-infected mice when compared to the dexamethasone alone (Fig. 8 and Table 4). In fact, despite the presence of some yeast, the *pdr6*Δ-infected lung tissue appears relatively healthy with little to no damage and inflammation suggesting that another mechanism may be responsible for the mice reaching endpoint (Fig. 8B and Table 4).

**Fig. 7.**
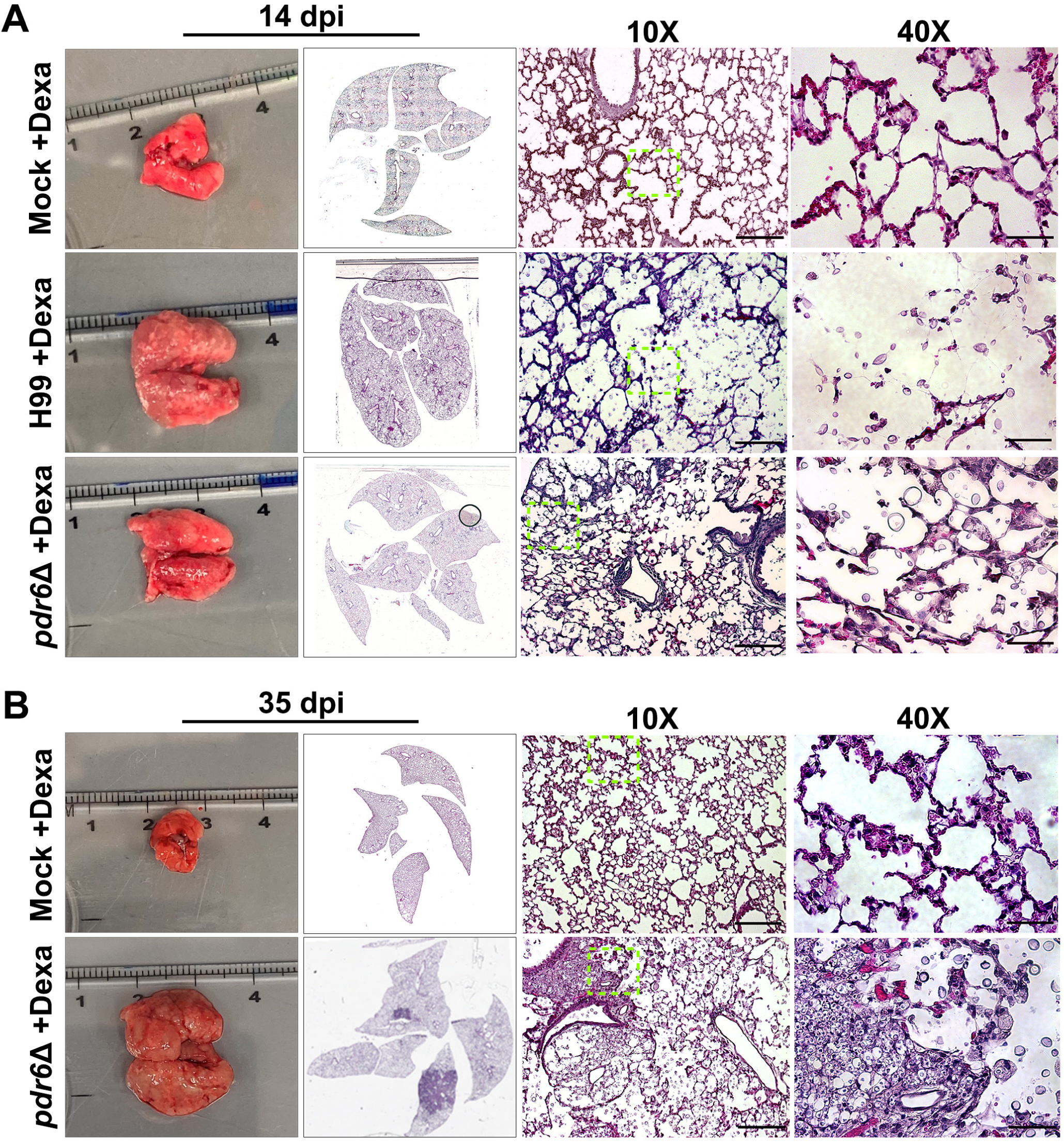
Effect of dexamethasone treatment on lung histopathology. Dissected lungs of female A/J mice infected with 5 x 10^4^ cells of the WT strain (H99) or *pdr6*Δ mutant strain and treated daily with 0.3 mg/kg Dexa and harvested at 14 and 35 dpi. Gross organ examination revealed alterations in levels of inflammation. H&E staining was used to visualize microscopic lung pathology (inset [green boxes]). The 10X scale bar is 200 μm and the 40X scale bar is 50 μm.

**Fig. 8.**
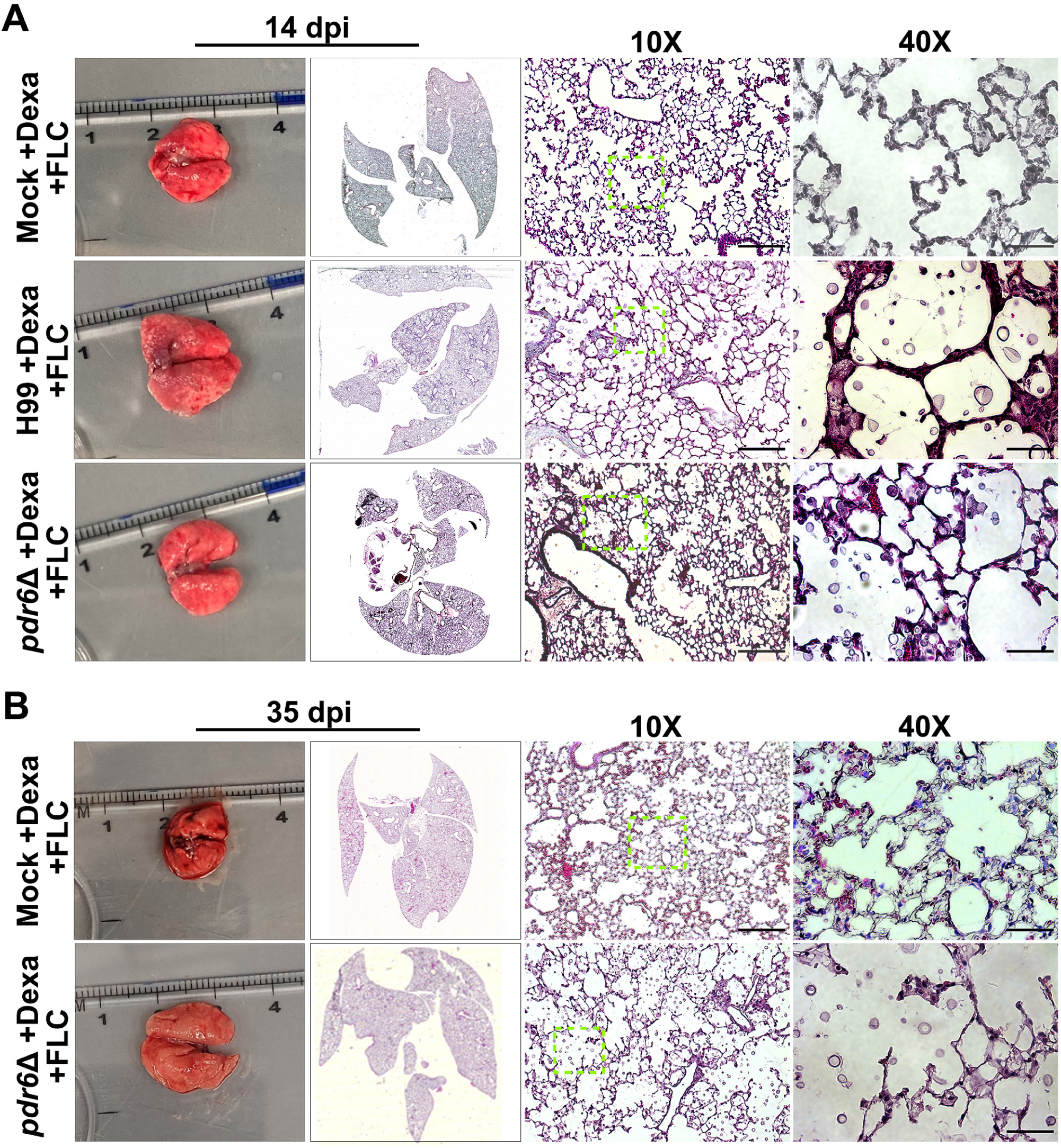
Effect of dexamethasone/FLC combination treatment on lung histopathology. Dissected lungs of female A/J mice infected with 5 x 10^4^ cells of the WT strain (H99) or *pdr6*Δ mutant strain and treated daily with a combination of 0.3 mg/kg Dexa and 16 mg/kg FLC and harvested at 14 and 35 dpi. Gross organ examination revealed alterations in levels of inflammation. H&E staining was used to visualize microscopic lung pathology (inset [green boxes]). The 10X scale bar is 200 μm and the 40X scale bar is 50 μm.

**Table 3.**
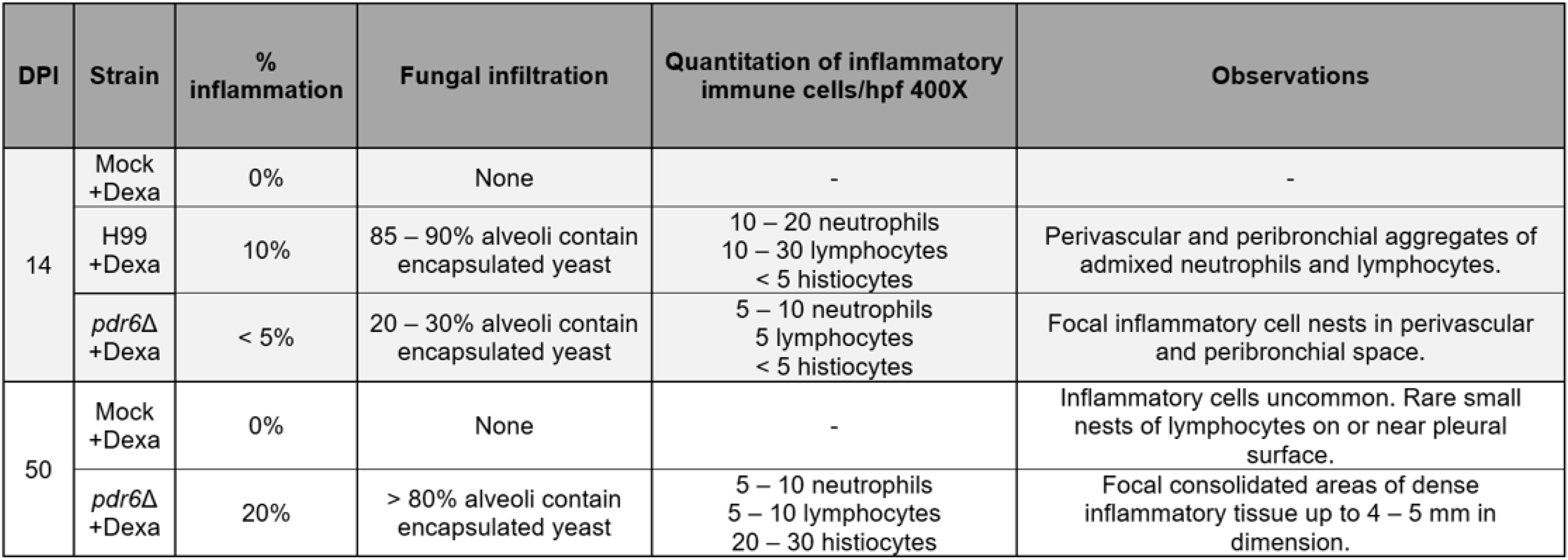
Quantitative analysis of dexamethasone-treated H99 and *pdr6*Δ-infected lung sections over time. Histiocytes refer to leukocytes of the monocyte/macrophage/dendritic cell lineage. hpf, high power field.

| DPI | Strain | % inflammation | Fungal infiltration | Quantitation of inflammatory immune cells/hpf 400X | Observations |
| --- | --- | --- | --- | --- | --- |
| 14 | Mock +Dexa | 0% | None | - | - |
|  | H99 +Dexa | 10% | 85 – 90% alveoli contain encapsulated yeast | 10 – 20 neutrophils<br>10 – 30 lymphocytes<br>< 5 histiocytes | Perivascular and peribronchial aggregates of admixed neutrophils and lymphocytes. |
|  | *pdr6Δ* +Dexa | < 5% | 20 – 30% alveoli contain encapsulated yeast | 5 – 10 neutrophils<br>5 lymphocytes<br>< 5 histiocytes | Focal inflammatory cell nests in perivascular and peribronchial space. |
| 50 | Mock +Dexa | 0% | None | - | Inflammatory cells uncommon. Rare small nests of lymphocytes on or near pleural surface. |
|  | *pdr6Δ* +Dexa | 20% | > 80% alveoli contain encapsulated yeast | 5 – 10 neutrophils<br>5 – 10 lymphocytes<br>20 – 30 histiocytes | Focal consolidated areas of dense inflammatory tissue up to 4 – 5 mm in dimension. |

**Table 4.** Quantitative analysis of combination Dexamethasone + Fluconazole treated H99 and *pdr6*Δ-infected lung sections over time. Histiocytes refer to leukocytes of the monocyte/macrophage/dendritic cell lineage. hpf, high power field.

| DPI | Strain | % inflammation | Fungal infiltration | Quantitation of inflammatory immune cells/hpf 400X | Observations |
| --- | --- | --- | --- | --- | --- |
| 14 | Mock<br>+Dexa<br>+FLC | 0% | None | - | - |
|  | H99<br>+Dexa<br>+FLC | 15 – 20% | 45 – 50% alveoli contain encapsulated yeast | 15 – 30 neutrophils<br>15 – 30 lymphocytes<br>< 5 histiocytes | Small to medium sized nests of inflammatory cells. |
|  | *pdr6Δ*<br>+Dexa<br>+FLC | 0% | 10 – 15% alveoli contain encapsulated yeast | - | Inflammatory cells are uncommon with rare perivascular and peribronchial nests of small lymphocytes and few histiocytes. |
| 50 | Mock<br>+Dexa<br>+FLC | 0% | None | Rare clusters of 2 – 4 histiocytes | Mildly dilated capillaries. |
|  | *pdr6Δ*<br>+Dexa<br>+FLC | 0% | 75 – 80% alveoli contain encapsulated yeast | - | Few small foci of perivascular lymphocytes and rare histiocytes. |

### *pdr6*Δ-infected mice display an increase in innate immune cells populations

During microbial infections, the cells of the innate immune system represent the first line of defense which is followed by the activation and recruitment of the adaptive immune response (31). In a *C. neoformans* infection, lung macrophages represent the innate population primarily responsible for initial clearance, although some recent literature suggests neutrophils also play a supportive role (32–34). However, CME is an AIDS-defining disease, and low CD4^+^ T-cell counts is a major susceptibility for CME, indicating that the adaptive immune response also plays an important role controlling cryptococcal infections (35). Provided the altered immune response to the *pdr6*Δ strain, we expected alterations to both the quantity and the composition of innate and adaptive inflammatory immune cells. To test this, we harvested leukocytes from the lungs of H99 and *pdr6*Δ-infected mice at 14 dpi (close to TOD of WT-infected mice) and at 35 dpi (close to TOD of *pdr6*Δ-infected mice) and quantified the different populations using flow cytometry (Fig. 9 and Fig. S2). Quantification of the total innate immune cell populations at 14 dpi showed that the FLC, dexamethasone, and combination treatments led to a significant decrease in the *pdr6*Δ-infected animals whereas there were no significant changes in the H99-infected mice (Fig. 9A). However, no reduction was observed when compared as a percentage of CD45^+^ cells (Fig. 9B), although, interestingly, a significant increase in the ratio of neutrophils and macrophages was seen in the combination-treated *pdr6*Δ-infected mice (Fig. 9B). At this time (14 dpi), quantification of the total adaptive immune cells showed a marked increase in the *pdr6*Δ-infected animals when compared to their H99-infected counterparts, although only the CD8^+^ T-cells were statistically significant (Fig. 9C, untreated bars). Furthermore, all treatments decrease the number of lymphocytes in the *pdr6*Δ-infected animals relative to the WT, specially the CD8^+^ T-cells (Fig. 9C, +FLC, +Dexa, and +Dexa +FLC bars). The same trend is observed when compared as percentage of CD45+ cells (Fig. 9D). At 35 dpi there are some minor differences in the adaptive immune cell populations when comparing the treated groups (Fig. 9G & H). Interestingly, when comparing the untreated *pdr6*Δ-infected mice at 35 dpi with 14 dpi there is a significant reduction in the total number of innate immune cells (Fig. 9D & H, untreated bars). Moreover, a significant increase in macrophages is observed in the *pdr6*Δ-infected mice when compared to the H99-infected (Fig. 9E & F, untreated bars). Using CD68 as a marker of inflammatory macrophages in histological slices confirmed this influx of immune cells specific for the *pdr6*Δ-infected (Fig. S3). Notably, the dexamethasone treatment (alone or with FLC) in the *pdr6*Δ-infected animals significantly decreases the number of macrophages which is replaced by a significant increase in the number of neutrophils (Fig. 9E & F, +Dexa and +Dexa +FLC bars). It appears that in the absence of Pdr6, a robust and sustained macrophage response is mediating the tissue damage, whereas in the WT-infected mice, neutrophils are the main innate cells accumulating in the lungs. Collectively, these results showed that infection with the *pdr6*Δ strain results in the increased recruitment of adaptive immune cells early on during the infection while an increase in innate immune cells, specifically macrophages, was observed in the later stages of the disease. This further explains the strong pro-inflammatory response, altered disease progression, and differential symptomatology specific to the *pdr6*Δ strain.

**Fig. 9.**
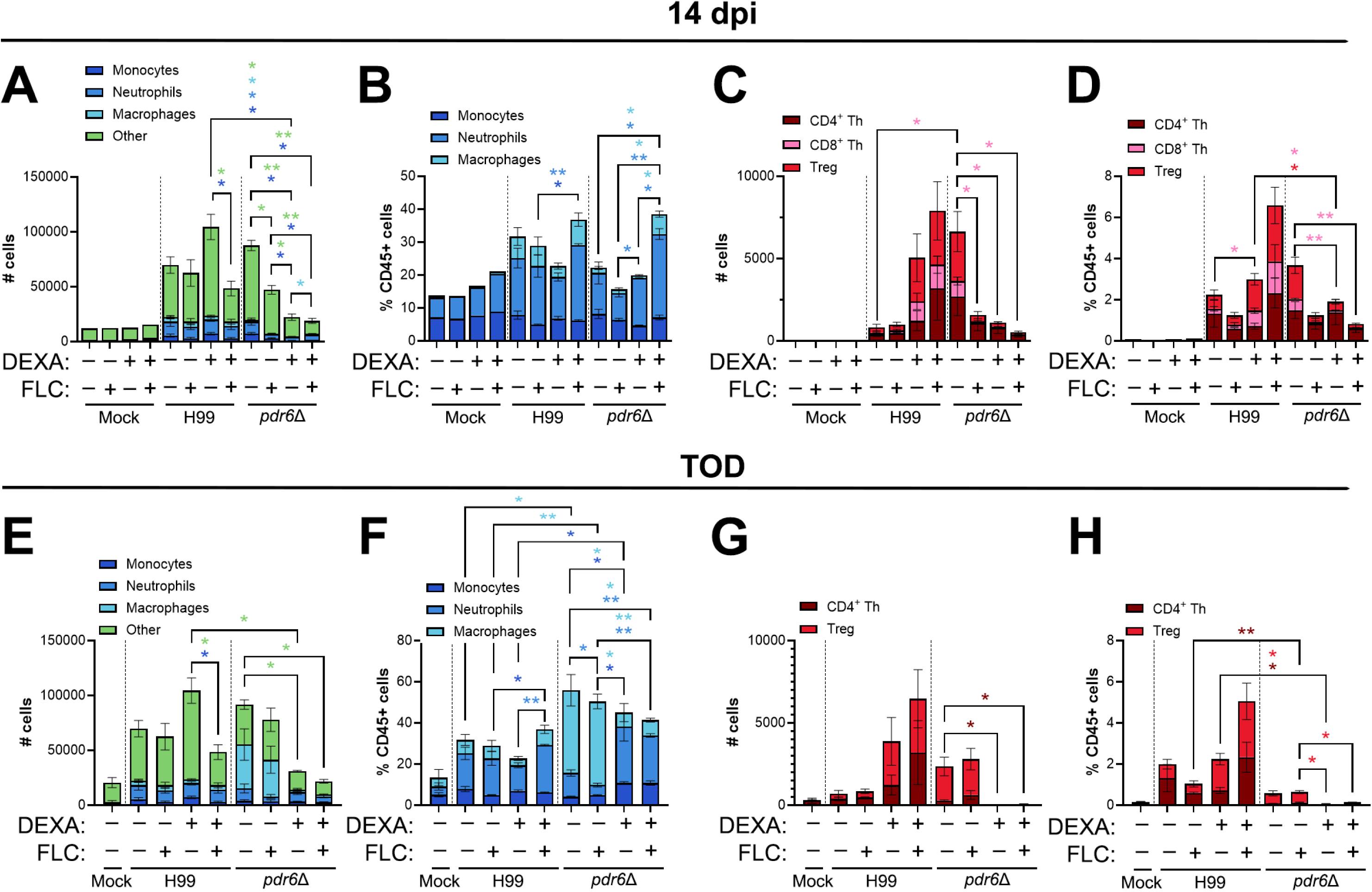
Infection with the *C. neoformans pdr6Δ* mutant generates an increase in innate immune cells, specifically macrophages. **(A-H**) Quantification of innate and adaptive immune cell populations at 14 dpi **(A-D)** and endpoint (35 dpi for *pdr6*Δ-infected mice)**(E-H)**. Female A/J mice were intranasally infected with 5 x 10^4^ cells of the WT type (H99) or *pdr6*Δ strain and euthanized at clinical endpoints corresponding to imminent mortality. Lungs were harvested and single-cell suspensions were generated and stained with immune panels to quantify monocytes (CD45+F4/80-CD11b+Ly6C+), neutrophils (CD45+F4/80-CD11b+Ly6G+), macrophages (CD45+F4/80+CD11b+), CD4^+^ Th (CD45+CD3+CD4+), CD4^+^ T-regs (CD45+CD3+CD4+CD25+FoxP3+), and CD8^+^ (CD45+CD3+CD8+) cells. Samples were processed using flow cytometry and are presented as total # cells **(A, C, E, G)** or as % CD45^+^ cells **(B, D, F, H)**. Significance was determined using one-way ANOVA with multiple comparisons (Welch’s corrected) *P<0.05; **P<0.01. Note: for **G** and **H**, the CD8^+^ T cell panel could not be run for the day 35 samples, hence they were removed from the H99 bars.

### The magnitude of the host response to *pdr6Δ* strain infection influences the disease outcome

Up to this point, our results indicate that infection with the *pdr6*Δ strain results in the recruitment of more inflammatory immune cells to the lungs, that initially can control the infection but, when sustained over several weeks, leads to increased pulmonary damage and mortality. This is supported by the dexamethasone treatment experiments, where dampening the immune response significantly extended survival in the *pdr6*Δ-infected animals. However, these experiments were done in the A/J mouse background, which is known to be “muted” in response to cryptococcal infection. We wondered if the same would happen in mice that are known to mount either strong pro-inflammatory (C57BL/6) or anti-inflammatory (BALB/c) responses against WT *C. neoformans*. With C57BL/6 mice, we envisioned two potential outcomes: the already skewed pro-inflammatory response of this mouse in combination with the pro-inflammatory response induced by *pdr6*Δ could result in complete clearance, or augment the tissue damage and result in hypervirulence. With the BALB/c mice, we had one hypothesis: the strong anti-inflammatory response of this mouse could counter the pro-inflammatory response induced by *pdr6*Δ enough to reach a balance, allowing for clearance or latency. To test this, we infected either C57BL/6 or BALB/c mice with 5 x 10^4^ cells of H99 or *pdr6*Δ strains and followed their survival as before. Whenever a mouse reached humane endpoints, organs were harvested for CFU and cytokine quantification. Interestingly, the C57BL/6 survival curve was similar to the A/J curve (Fig. 10A), with a median survival time of 18 days for H99-infected mice and 42 days for *pdr6*Δ-infected mice, and decreased *pdr6*Δ burden in all organs (Fig. 10B – D). Indeed, the cytokine profile of the *pdr6*Δ-infected C57BL/6 was a more pronounced pro-inflammatory A/J profile, with the same upregulation of IL-1α, IL-1β, and MIP-1β, but additionally, there was strong upregulation of IL-6, MIP-1α, and G-CSF (Fig. 10E and Fig. S4). In contrast, 70% of the *pdr6*Δ-infected BALB/c mice survived to the end of the experiment (60 dpi), whereas all of the H99-infected mice succumbed by day 18 (Fig. 10F). Surprisingly, the surviving mice had fungal burdens higher than the H99-infected mice at TOD (Fig. 10G – I), yet they showed no symptoms. Comparison of the immune profiles between C57BL/6 and BALB/c showed expected similarities (IL-1α, IL-1β, and MIP-1α) but also clear differences (lack of IL-6 and G-CSF signaling and stronger induction of IL-13 in BALB/c)(Fig. 10E & J, and Fig. S4). Similarly to the dexamethasone treatment in A/J mice, the *pdr6*Δ-infected BALB/c mice survived longer despite having higher fungal burdens, strongly suggesting that the fatal pathology is driven by the immune response and not by the fungus.

**Fig. 10.**
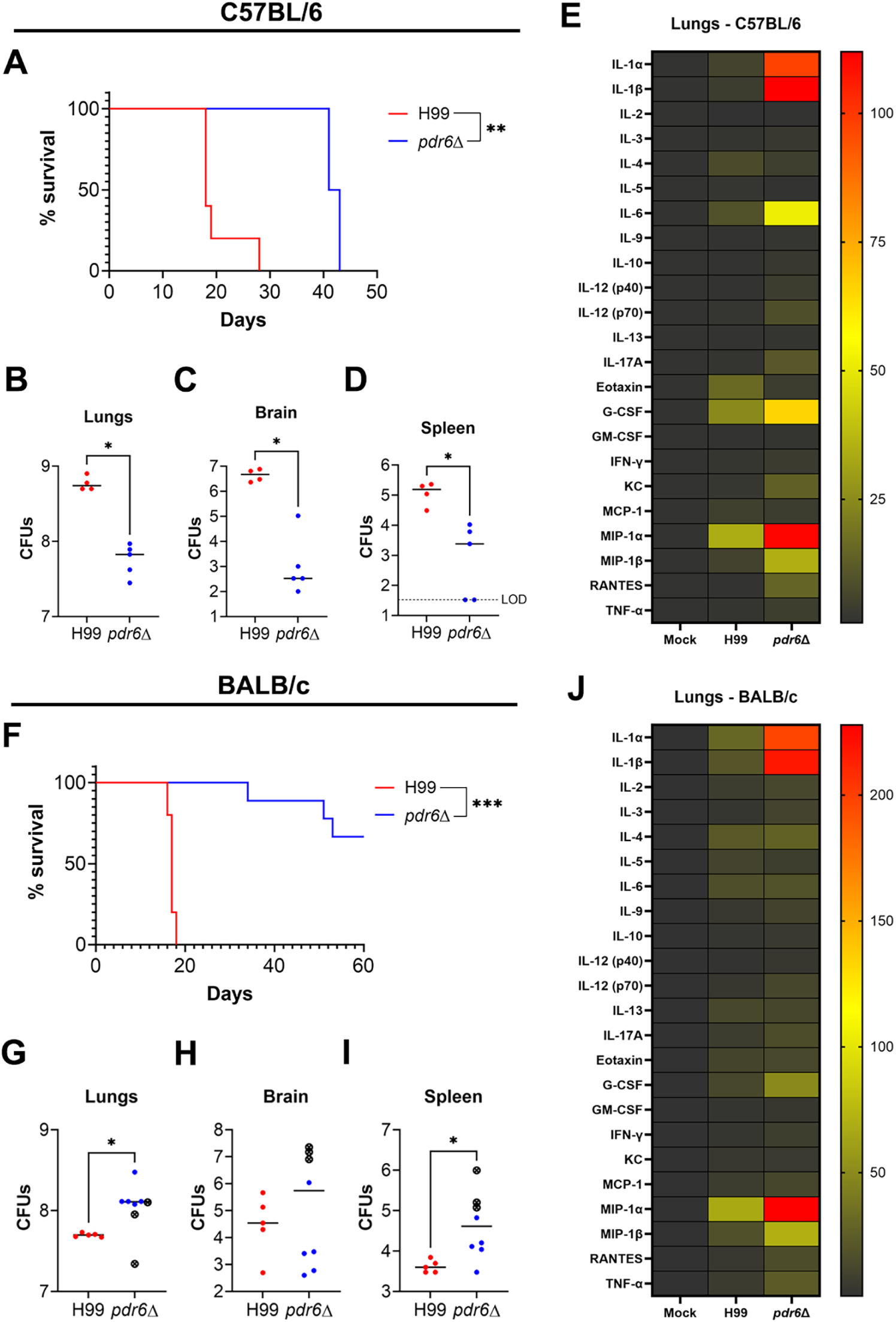
Mice breeds with skewed immune responses influence the outcome of the *pdr6*Δ infection. Survival curves and organ burdens for C57BL/6 **(A – D)** or BALB/c **(F – I)** mice infected with 5 x 10^4^ cells of WT (H99) or *pdr6*Δ strains. At least 5 mice were infected for each group. Significance in the survival curves was determined using Mantel-Cox test, **P<0.01; ***P<0.001. For fungal burdens, organs were harvested, homogenized, and dilutions were plated on YPD. Plates were incubated for 48 hr and CFUs were enumerated. Each data point represents a mouse and black bars represent the mean. In panels G-I, crossed data points represent the values from the mice that succumbed to infection. Significance was determined using one-way ANOVA with multiple comparisons (Welch’s corrected), *P<0.05. **(E & J)** Heatmap of the pro-inflammatory and anti-inflammatory cytokines/chemokines in the lungs of C57BL/6 **(E)** and BALB/c **(J)** infected mice. Values are shown normalized to age-match mock-infected mice.

### *C. neoformans pdr6Δ* strain exhibits altered PAMP exposure

Experimentally, we have studied the host response to this cryptococcal mutant but the question remains as to how is this mutant modulating the immune system and generating a robust Th1 response. In our previous work, we showed that *PDR6* encodes an atypical transporter localized to the endoplasmic reticulum (ER) and plasma membrane (PM) involved in ergosterol trafficking (13). Others have suggested a direct relationship between altered ergosterol levels and host immune responses (36, 37) as well as altered surface composition due to defects in PM fluidity and lipid rafts (38, 39). We hypothesize that in the absence of *PDR6* there would be less ergosterol in the PM, potentially resulting in downstream alterations to the cryptococcal surface that would expose immunogenic components. To assess this, we grew H99 and *pdr6*Δ cells in YPD, stained with fluorescent dyes that recognize and bind to fungal cell wall components, and visualized the cells using microscopy. Under this condition, we did not observe any difference between the H99 and *pdr6*Δ stained cells (data not shown). However, previous studies have reported a correlation between growth conditions and the architecture of the fungal cell wall (40). This led us to repeat the experiment with cells grown under host-like conditions (DMEM at 37°C + 5% CO_2_) (Fig. 11A). Under these conditions, the *pdr6*Δ strain displayed increase staining with calcofluor white (CFW) and concanavalin A (ConA) suggestive of increased exposure of chitin and mannoproteins, respectively. Similarly, there was some staining with the β-(1,3)-glucan antibody suggestive of more exposed β-glucans in the *pdr6*Δ mutant (Fig. 11A). Despite the differences observed microscopically, quantification of the staining intensity of these dyes, and of eosin Y (EoY) which stains chitosan, using flow cytometry did not reveal statistically significant increases in intensity in the *pdr6*Δ strain when compared to the H99 (Fig. 11B – E). The positive control, *cap59*Δ, deficient in polysaccharide capsule synthesis and thus with increased exposure of its cell wall, showed the expected increases in intensity. This was confirmed using three independently made *pdr6*Δ mutants (Fig. 11B – E and Fig. S5). Having previously shown that the *pdr6*Δ mutant has no defect in capsule induction, we hypothesized that alterations in the capsular architecture could explain the visually increased exposed cryptococcal PAMPs. To assess this, we used several anti-polysaccharide capsule antibodies to stain the capsules of H99, *pdr6Δ*, and *cap59*Δ cells (Fig. 11F). A significant increase in staining and penetration of the antibody was observed in the *pdr6*Δ mutant when using the mAb 302 (Fig. 11F). This antibody only recognizes and binds acetylated GXM (41) and suggests that the integrity and/or porousness of the polysaccharide capsule in the *pdr6*Δ strain is altered when compared to the structurally-sound H99 capsule. This could justify the high-uptake phenotype of the *pdr6*Δ mutant (13, 42). Lastly, we tested if the *pdr6*Δ mutant could form titan cells using a recent *in vitro* induction protocol (43), as titan cells are restricted to the lungs, have an altered cell surface, and are related to latency. Although there was a trend to form less titan cells, it was not statistically significant relative to H99 (Fig. 11G). Altogether, these findings suggest that the deletion of *PDR6* disrupts the cryptococcal cell wall composition and the capsule density (since it is induced to normal size in the *pdr6*Δ mutant), increasing the potential exposure of fungal PAMPs, which can drive the strong pro-inflammatory response by the host immune system.

**Fig. 11.**
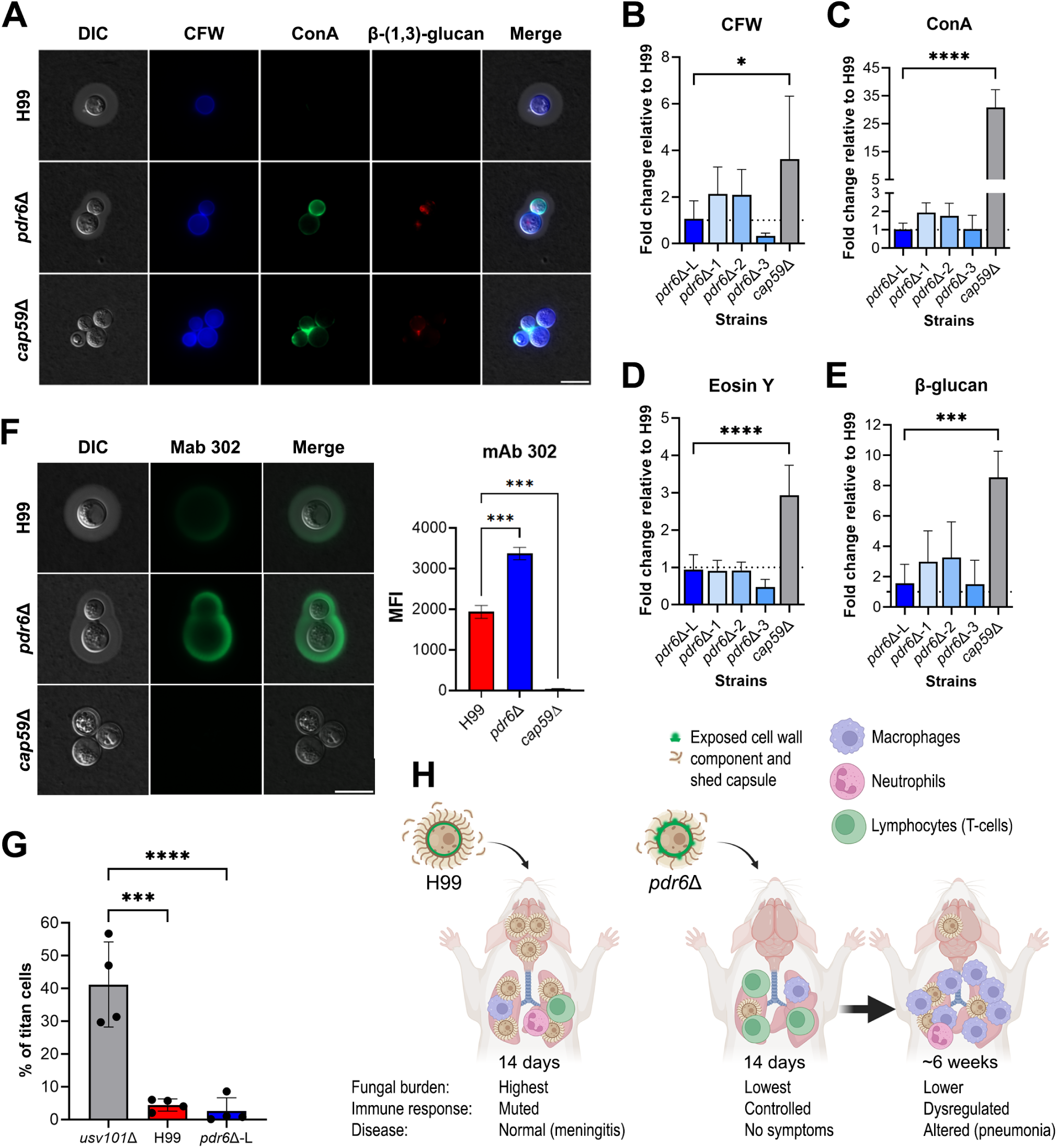
*C. neoformans pdr6*Δ cells display altered cell wall features and capsule density. **(A)** Visualization of calcofluor (CFW), Alexa488-conjugated Concanavalin A (ConA), and β-(1,3)-glucan staining in H99, *pdr6*Δ-L (from commercial deletion collection), and *cap59*Δ cells. Representative images of stained cells. Scale bar, 10 μm. (B - E) Quantification of CFW (B), ConA (C), Eosin Y (D), and β-glucan **(E)** staining intensity in H99, *pdr6*Δ, and *cap59*Δ strains. Staining intensity was measured by flow cytometry and results normalized to H99 (dotted line). Four independent *pdr6*Δ strains were tested (*pdr6Δ*-L and *pdr6*Δ-1 to -3). Significance was determined using one-way ANOVA (Welch’s corrected), *P<0.05; ***P<0.001; ****P<0.0001. **(F)** Visualization and quantification of capsular antibody staining of H99, *pdr6*Δ, and *cap59*Δ strains. Cells were stained with anti-capsule mAb 302 and imaged using an inverted Zeiss microscope. Representative images of stained cells. Staining intensity was quantified using a Cell Profiler pipeline. Significance was determined using one-way ANOVA (Welch’s corrected) ***P<0.001. **(G)** The % of titanization of the indicated strains was determined as described in the methods. Each data point represents the average % of titanization in independent biological experiments each with at least 100 cells quantified. One-way ANOVA with multiple comparisons, ***P<0.001; ****P<0.0001. **(H)** Model explaining how *pdr6*Δ strains alter disease progression in A/J and C57BL/6 mice, causing fatal pneumonia rather than the typical meningitis caused by H99. The *pdr6*Δ strain has an altered cell surface, including a less dense capsule that is more permeable to antibodies, that results in exposure of pro-inflammatory antigens. This results in an initial immune response that can control the infection, consisting on an increased number of CD4 lymphocytes. However, that initial response is not enough to clear the infection, and eventually results in dysregulation, with a drop in the number of CD4 lymphocytes and a dramatic increase in inflammatory macrophages. Despite having less fungal burden in the lungs and little dissemination, the mice succumb to the infection due to too much lung tissue damage. This illustration was originally made in BioRender https://BioRender.com/vpa8dnh) and subsequently edited in Photoshop.

## DISCUSSION

Fungal PDR transporters have been primarily studied for their role in antifungal resistance. For example, the overexpression of *AFR1* in *C. neoformans* and *CDR1* in *C. albicans* clinical strains increase their resistance to many of the azole class antifungals and complicates their treatment strategy (44, 45). Structurally, these two proteins (and all other well-characterized fungal PDR transporters) are ‘full-length’ consisting of two nucleotide-binding domains (NBDs) and two transmembrane domains (TMDs), which is required for a functional fungal PDR protein (15, 16, 46). Here, we continue the characterization of *C. neoformans PDR6*, an atypical and novel ‘half-size’ PDR transporter since it only possesses one NBD and one TMD. Despite this, various lines of evidence suggest it forms a functional transporter (13, 14). However, we found that in addition to azole resistance, *PDR6* modulates the host immune response, a phenotype that may be unique to ‘half-size’ PDR transporters (13).

The *pdr6*Δ-dependent alterations in survival, fungal burden, and symptoms all suggested a differential host response. Moreover, the fact that despite the significant decrease in fungal burden across organs, and the use of a lower inoculum, the mice still died, suggesting that the *pdr6*Δ-infected mice were not succumbing to uncontrolled fungal growth, but rather to an inappropriate and damaging immune response. This was further supported by the FLC and dexamethasone studies. Treatment of the *pdr6*Δ-infected mice with FLC decreased the fungal burden in all organs, and in some to the limit of detection, and yet the mice still died. In contrast, but consistent with the immune response as the culprit, dexamethasone treatment in the *pdr6*Δ-infected mice significantly extended their survival despite having fungal burdens that were higher than the WT-infected mice at TOD. Lastly, the divergence in symptoms, histological analysis, immune cell quantification, and cytokine/chemokine expression profiles, all were consistent with an aggressive lung pro-inflammatory immune response against the *pdr6*Δ mutant, resulting in their death by pneumonia rather than by CME. We recognize, however, that the *pdr6*Δ-infected mice still had CFUs in the brain, so CME cannot be ruled out, but we believe that it was below the threshold to trigger the neurological symptoms characteristic of CME. This is supported by the “muted” immune response in the brain tissue, indicative of little inflammation, and by the fact that when treated with FLC, there was minimal extrapulmonary dissemination but no change in the outcome of the infection. If we consider that the *pdr6*Δ mutant grows slower under host-like conditions (13), is recognized more avidly by THP-1 cells (alveolar macrophages)(13, 42), and has reduced intracellular survival (13, 14), then the massive lung tissue damage seen in the *pdr6*Δ-infected mice at TOD has to be due to an inappropriate, dysregulated, Th1 immune response.

Although treatment with a corticosteroid supported our hypothesis, we acknowledge there are some limitations. Dexamethasone is a strong immunosuppressant, dampening the immune response in the whole body, especially given the length of the study. To address that, we purposefully used doses that corresponded to less than those used in human clinical trials (29). This extended treatment might explain why, even in the combo treatment with FLC, which reduced both inflammation and fungal burden in the lungs and restored tissue architecture almost to normal, the mice appeared so sickly. Optimizing the timing of the dexamethasone treatments, or employing ways to target the drug specifically to the lungs, as with the use of an inhaler, might results in improved outcomes and represents a future direction of investigation. However, we reasoned that we could counter the pro-inflammatory response induced by the *pdr6*Δ mutant using mouse breeds that naturally mount anti-inflammatory, or Th2-skewed, responses to infection, such as the BALB/c mouse. As a control, we also infected C57BL/6 mice, which are known to be Th1-skewed in their immune responses. Again, consistent with our hypothesis, 70% of the BALB/c mice survived until the experimental endpoint at 60 dpi despite having significantly higher lung fungal burdens at this timepoint compared to the H99-infected mice at TOD.

The *in vivo* experimentation allowed us to comprehend what was occurring within the host, but why was this specifically occurring with the *pdr6*Δ strain? Our data showing that the *pdr6*Δ mutant displays altered PAMP exposure is a reasonable explanation for the pro-inflammatory response. However, it may not be the only reason as the acapsular *cap59*Δ mutant also exhibits increased PAMP exposure and it is avirulent in the same mouse model. Furthermore, we previously showed that conditioned media (CM) from *pdr6*Δ cultures modulates the phagocytosis of *C. neoformans* by macrophages, whereas the CM from *cap59*Δ or other PDR transporter mutants had little to no effect (13), suggestive of an immunomodulatory Pdr6-dependent secreted factor. Furthermore, other cryptococcal mutants with altered PAMP exposure have been reported but none of them induce an immune response as that seen here with the *pdr6*Δ mutant (47–50). The *mar1*Δ mutant has a disorganized cell wall with increased chitin exposure, less glucans and mannans, and little capsule (49). Like our case, the *mar1*Δ mutant causes a protracted death in C57BL/6 but shows no mortality in BALB/c (47). However, in the *mar1*Δ-infected BALB/c mice, the immune response was controlled in the form of GM-CSF-dependent granulomas and there was no overt pro-inflammatory response, which is different from *pdr6*Δ-infected mice. The *pdr6*Δ-induced mortality is more similar to that seen with the *usv101*Δ strain, which causes fatal pneumonia in A/J mice with little to no extrapulmonary dissemination (50). However, pathologically, the *usv101*Δ mutant did not generate granulomas like the *pdr6*Δ mutant, and in our case, we see much more extrapulmonary dissemination than that with the *usv101*Δ mutant. The *usv101*Δ strain has a larger capsule than WT, thus it might not be exposing the same antigens as the *pdr6*Δ (which has a normal-sized capsule), generating a muted immune response relative to the response induced by *pdr6*Δ. Lastly, the *rim101*Δ mutant also induces a strong pro-inflammatory response in A/J mice as a result of an altered cell surface, but unlike the *pdr6*Δ mutant, it results in hypervirulence rather than a protracted death. Furthermore, the *rim101*Δ-induced immune response is mediated by neutrophils and eosinophils, hence, does not involve granulomas like in the *mar1*Δ or *pdr6*Δ cases. Therefore, an altered exposure of PAMPs alone cannot explain the *pdr6*Δ modulation of the immune response. Consistently, there are examples of cryptococcal mutants that induce altered immune responses without any detectable surface changes, like the F-box protein mutant *fbp1*Δ (48). Overall, this work and the examples discussed indicate that a granulomatous inflammatory response is protective against *Cryptococcus* lung infection. Interestingly, early clinical studies in cryptococcosis patients showed that T cells are needed for an effective containment of fungal cells in a granuloma (51). This is consistent with our finding that at 14 dpi, when the mice are controlling the *pdr6*Δ infection, there is a considerably increase in lymphocytes relative to the WT-infection, while there are no significant changes in innate cells. However, at 35 dpi, percentage-wise, there are almost no lymphocytes in the *pdr6*Δ-infected lungs while there is a significant increase in macrophages (Fig. 11H). Although further studies depleting these cell populations are needed to confirm this, it is conceivable that a failed sustained lymphocyte response is responsible for the eventual death in the *pdr6*Δ-infected mice. Curiously, a recent study from the Nielsen lab found an association between hypervirulence and increased Th1/Th17 immune responses specific to the A/J mouse breed (52). However, in that study, the hypervirulent strains induced a neutrophil-based response while the strong inflammatory response induced by *pdr6*Δ seems to be mediated by macrophages and not by neutrophils (Fig. 11H).

Taken together, these data provide important insights for improving the management of CME. Anticryptococcal treatments are limited by toxicity, costs, and rising microbial resistance, but FLC, despite being sub-optimal, still is the antifungal of choice in many regions of the world because it is cheap, relatively less toxic, and broad spectrum. Since the deletion of *PDR6* results in hypersensitivity to FLC, the design and synthesis of a novel compound that targets Pdr6, coupled with FLC and/or a corticosteroid, may have clinical potential. Alternatively, several groups in the cryptococcal and immunology fields have tried to identify potential vaccine candidates against this fungus (53). Collectively, these preliminary vaccines, elicit a pro-inflammatory response allowing for protection against subsequent challenges (54–56). However, this is hampered because the immune mechanisms that elicit a protective immune response are not known yet. Thus, the knowledge generated in this study has potential to help inform cryptococcal vaccine development by providing novel insights into what constitutes a deleterious pro-inflammatory response against *C. neoformans* infection.

Finally, the full complexities of PDR half-size transporters remain unknown. It is understood that these proteins are promiscuous and transport a diverse array of substrates, and with the expanded repertoire of PDR genes in *Cryptococcus*, it is not surprising to find roles other than traditional efflux pumps and subsequent resistance to xenobiotics. *PDR6* has also been implicated in aging and replicative life span (14), and most recently, *PDR9*, a full-length PDR gene in the same clade as *AFR1*, was found to play a role in CO_2_ tolerance via an unknown mechanism that may involve phospholipid flipping (57). Hence, *PDR6* may act through multiple direct and indirect mechanisms to modulate the inflammatory host response, such as by secreting immunomodulatory molecules, aiding in maintaining the correct PM fluidity by regulating ergosterol transport, and preventing exposure of surface PAMPs by supporting correct cell wall and/or capsule architecture. All of these functions could be mediated by Pdr6-dependent metabolites, lipids, or proteins, supporting future studies in these areas such as comparative metabolomics of WT and *pdr6*Δ CM.

In conclusion, we have further characterized a novel PDR transporter in *C. neoformans* by studying the role of *PDR6* in a mouse model of cryptococcosis and examining the altered immune response. Mice infected with the *pdr6*Δ strain develop an overexuberant inflammatory response that is effective at controlling the infection in the lungs short-term, but eventually causes a fulminant pneumonia. Elucidating the role of *PDR6* in the context of infection improves our understanding of cryptococcal pathogenesis while also informing potential clinically-relevant therapeutic studies such as the search for novel drug targets or the development of a vaccine. More broadly, this study expands the field of fungal PDR transporters and given the conserved nature of these proteins, these findings may be applicable to other fungal pathogens.

## MATERIALS & METHODS

### Strains, media, and growth conditions

All *C. neoformans* strains used in this study were in the serotype A H99α background. All fungal strains were maintained at −80°C and grown at 30°C on yeast peptone dextrose (YPD) with antibiotics (NAT or G418) as appropriate. The *pdr6*Δ strain used for most of these studies was obtained from the Madhani deletion collection (58) and the recreated strains were made in this background by biolistics (13). The genome of all four independently created *pdr6*Δ mutants was sequenced to confirm proper deletion of the gene without additional gross changes.

The human monocyte cell line THP-1 (ATCC TIB-202) was grown in THP-1 complete medium and differentiated with phorbol 12-myristate 12-acetate (PMA, Sigma) as described (13). The hAELVi cell line is an immortalized normal human alveolar epithelial cell obtained commercially (InSCREENeX GmbH, Germany) and was grown and used as described (59).

### Murine survival studies

*In vivo* fungal virulence was tested using the murine inhalation model of cryptococcosis (13, 42, 50, 60). Briefly, 5- to 6-week-old female inbred mice (A/J, C57BL/6, or BALB/c, all from The Jackson Laboratories) were anesthetized with a ketamine rodent cocktail delivered i.p. and intranasally inoculated, unless otherwise stated, with 5.0 x 10^4^ cryptococcal cells in 40 μL of sterile PBS. Animals were closely monitored and humanely euthanized if they lost >20% weight relative to peak weight, or at predetermined timepoints, or at the end of the experiment (maximum of 60 days). For drug treatment studies, cryptococcal-infected mice were randomly assigned into treated or non-treated control groups. Stock solutions of fluconazole (1.6 mg/mL), dexamethasone (0.03 mg/mL), and a fluconazole/dexamethasone combo were prepared in sterile PBS. The drug(s) were administered intraperitoneally at doses of 16 mg/kg/day or 8 mg/kg/day for fluconazole, and of 0.3 mg/kg/day for dexamethasone. Treatment with these compounds began 24 h after infection and continued throughout the experiment until endpoint was reached. Statistically significant differences between survival curves were determined by log rank Mantel-Cox test (GraphPad Prism). All of the mouse infections and procedures were approved by the University of Notre Dame Institutional Animal Care and Use Committee (protocol #23-05-7918).

### Fungal Burden Quantification

Infected mice were humanely euthanized at predetermined time points or TOD by isoflurane overdose followed by cervical dislocation, and lungs, brain, and spleens were harvested. Tissues were homogenized in 5 mL of sterile PBS, diluted, and plated on YPD agar medium supplemented with ampicillin. CFUs were counted following incubation at 30°C for 48 h. Statistical significance was determined using ANOVA Kruskal-Wallis test (GraphPad Prism).

### Histological analyses

A/J female mice (The Jackson Laboratories) were infected via the intranasal route as described above with 5 x 10^4^ CFU of either WT (H99) or the *pdr6*Δ mutant strain. Mice were sacrificed as predetermined time points (14, 35, 50 days post inoculation) by isoflurane overdose followed by cervical dislocation. Lungs were perfused with sterile PBS and 10% neutralized formalin. Lungs were subsequently paraffin embedded, sectioned, mounted, and stained with H&E by the University of Notre Dame Histology Core facility. H&E-stained slides were imaged at 10X and 40X using an inverted Zeiss microscope. Slides were scored in a blind manner by the resident pathologist at the Harper Cancer Research Institute.

### Cytokine Analyses

Female mice (A/J, C57BL/6, or BALB/c, all from The Jackson Labs) were infected via the intranasal route with WT (H99) or the *pdr6*Δ mutant strain as described above. Mice were euthanized at predetermined time points by isoflurane overdose followed by cervical dislocation. Cytokine levels within the A/J mice lung and brain homogenates were analyzed using the Proteome Profiler Mouse XL Cytokine Array (R&D Systems Inc., Minneapolis, MN). The relative expression levels of 111 soluble mouse proteins were determined following the manufacturer’s protocol. Array membranes were imaged using LI-COR Odyssey Infrared Imaging System and quantified using ImageStudio software. Cytokine/chemokine expression levels were further confirmed in A/J mice, as well as in C57BL/6 and BALB/c mice, using the Bio-Plex Multi-Plex Assay Kit (Bio-Rad Laboratories, Hercules, CA). Supernatants from organ homogenates were used to determine presence of 22 proteins including eotaxin, G-CSF, GM-CSF, IFN-γ, IL-1α, IL-1β, IL-2, IL-3, IL-4, IL-6, IL-9, IL-10, IL-12 (p40), IL-12 (p70), IL-13, IL-17A, KC, MCP-1, MCP-1α, MIP-1β, RANTES, and TNF-α.

### Pulmonary leukocyte isolation and flow cytometric analysis

H99 and *pdr6*Δ-infected A/J mice were euthanized at predetermined time points and TOD by isoflurane overdose followed by cervical dislocation. Lungs of infected mice were excised and digested enzymatically at 37°C for 60 min in 10 mL of digest buffer (0.34 U/mL Liberase^TM^ (Roche) and 100 ug/mL DNaseI (ThermoFisher)) in 1X DPBS. After 60 min, digestion was arrested with 1X DPBS supplemented with 2% FBS and 0.2 um EDTA. The cell suspension was washed and passed through 40-μm cell strainers. The single cell suspensions were stained with a live/dead far red viability dye (Invitrogen), washed, and incubated on ice for 20 min with purified anti-mouse CD16/CD32 antibodies (BioLegend) to block Fc receptors. Following blocking, cells were incubated with fluorochrome-conjugated antibodies to allow for multistaining for 45 min on ice. The innate panel consisted of PE-Cy5 conjugated anti-CD45 (30–111), Brilliant Violet (BV) 650-conjugated anti-CD11b (M1/70), Super Bright (SB) 780-conjugated anti-F4/80 (BM8), BV711-conjugated anti-Ly6G (1A8), PE-Cy7-conjugated anti-Ly6C (HK1.4), PE-conjugated anti-SiglecF (1RNM44N), and APC-eFluor 780-conjugated anti-CD11c (N418). The adaptive panel included PE-Cy5 conjugated anti-CD45 (30–111), AF700-conjugated anti-CD3 (145-2C11), BV421-conjugated anti-CD4 (GK1.5), BV510-conjugated anti-CD25 (PC61-5), PE-conjugated anti-CD8 (53-6.7), and AF488-conjugated anti-FoxP3 (FJK-16s). Cells were washed and fixed with 2% formalin in FACS buffer. UltraComp eBeads (Invitrogen) were stained with each respective fluorochrome-conjugated antibody and used to determine and spillover/compensation calculations. Data was analyzed using FlowJo software.

### Cell wall and capsule stains

Starter cultures were grown overnight in YPD, washed with DMEM, and 1 x 10^6^ cells/mL were added to 24-well tissue culture plates. Plates were incubated at 37°C + 5% CO2 for 48 h. The cells were collected by centrifugation at 3,000 x g for 5 min. For staining, Calcofluor White (CFW), Alexa Fluor 488-conjugated concanavalin A (ConA), β-(1,3)-glucan antibody 2GB (ab233743; Abcam, USA), eosin Y (EoY), and the capsular antibody 302 were used. Cells were washed and stained with CFW, ConA, β-(1,3)-glucan antibody, EoY, and Mab 302 at 100 μg/mL, 50 μg/mL, 20 μg/mL, 500 μg/mL, and 10 μg/mL, respectively. Cells were incubated with the stains for 30 min, washed, fixed, and imaged at 100X using an inverted Zeiss microscope. Fluorescence of mAb 302 was calculated using Cell Profiler software. Flow cytometry data was acquired using a BD LSRFortessa cytometer. The CFW signal was detected using a 405 nm laser and the ConA and EoY signals were detected using a 488 nm laser. Flow cytometry data was analyzed with FlowJo software.

## ACKNOWLEDGMENTS

We acknowledge assistance and support on the murine studies from the University of Notre Dame Freimann Life Science Center (FLSC). We give a special thanks to William J. Kaliney, M.D., pathologist from Harper Cancer Research Center, for assistance with histological analysis and scoring. We acknowledge financial support from the Warren Center for Drug Discovery (pilot grant) and NIH grant R21AI71742. Other related work in the Santiago-Tirado lab is supported by NIH grant R01AI177875. We also thank members of the Santiago-Tirado lab for their thoughtful comments and feedback on this work.

---

**Figure S1.**
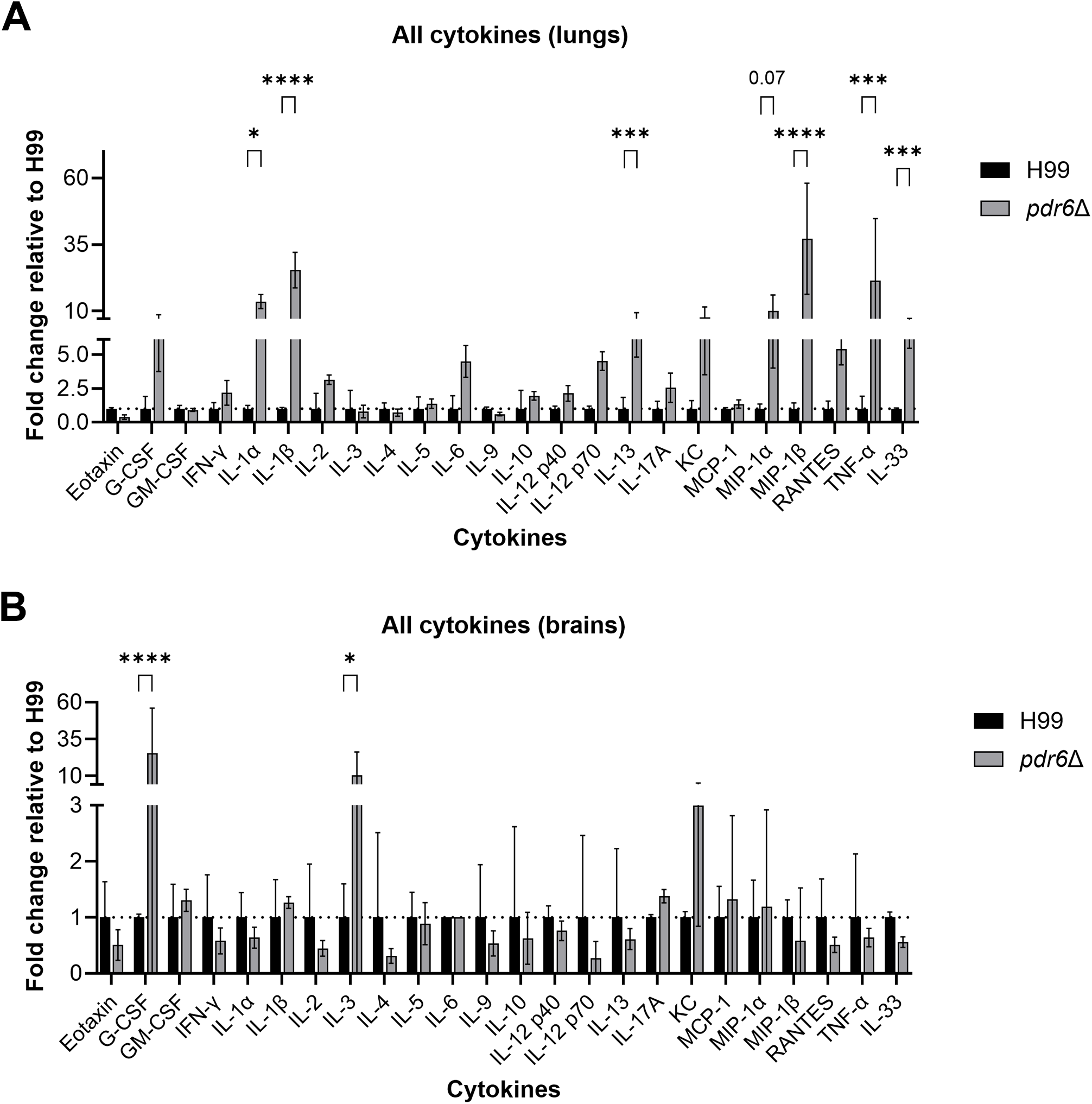

**Figure S2.**
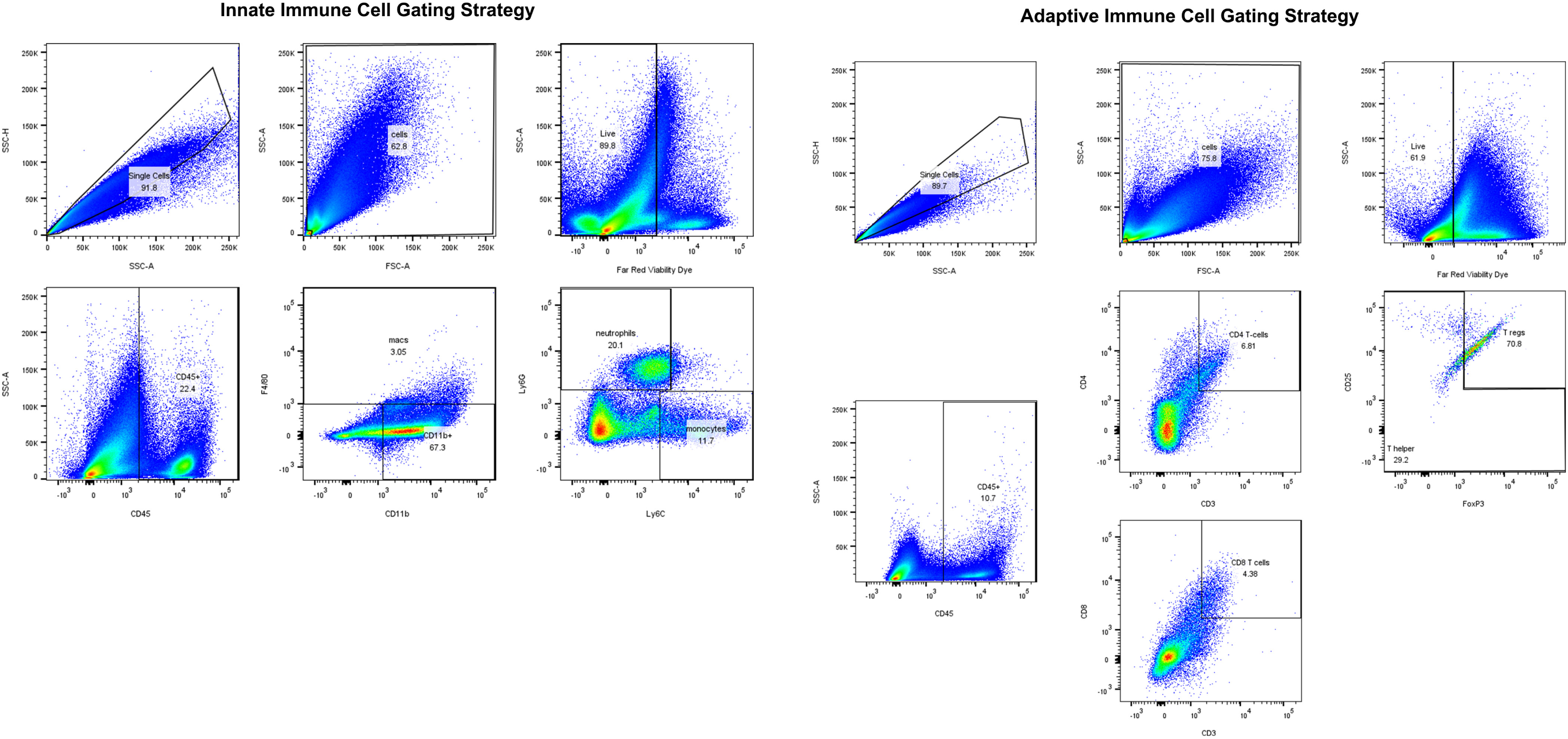

**Figure S3.**
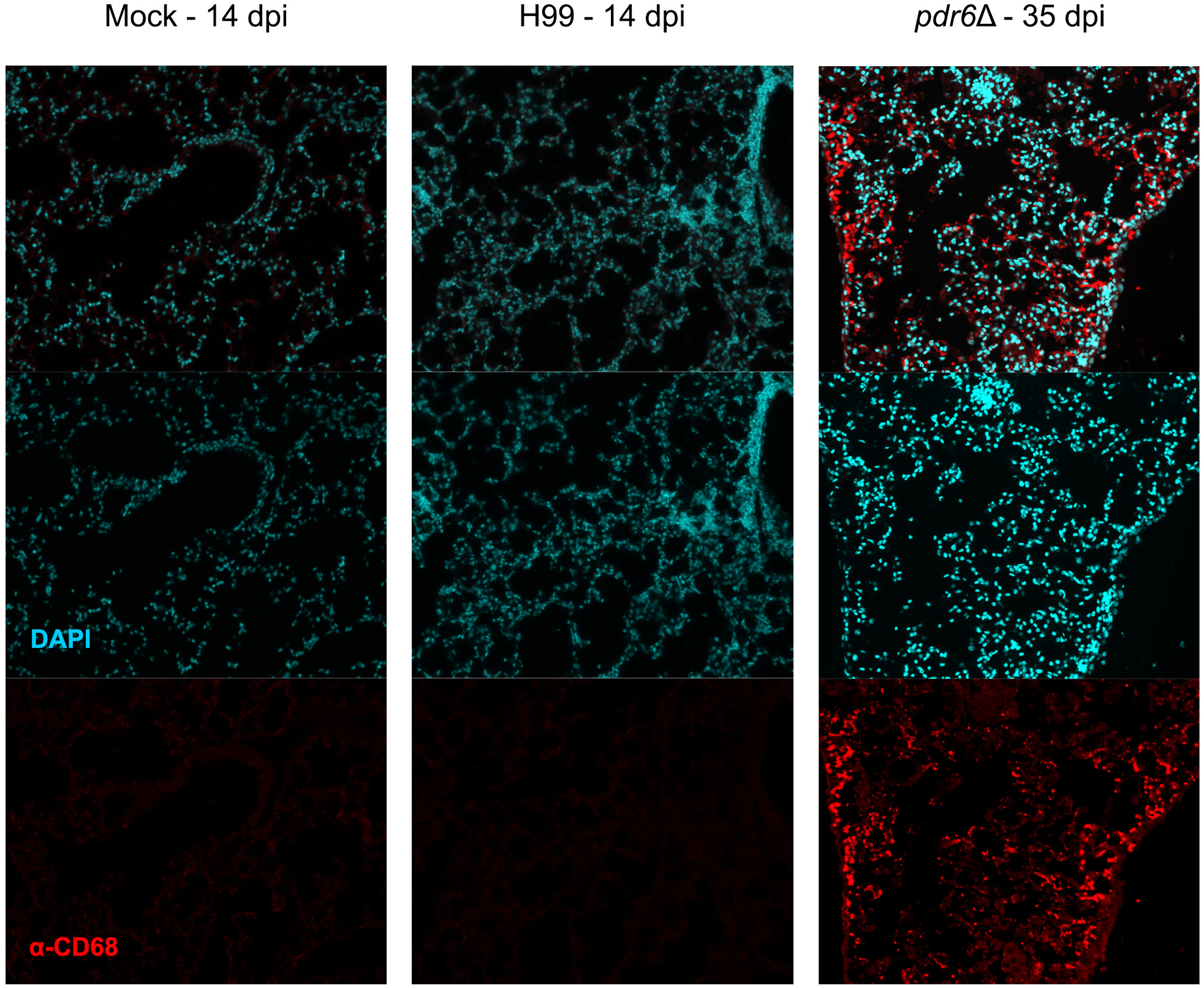

**Figure S4.**
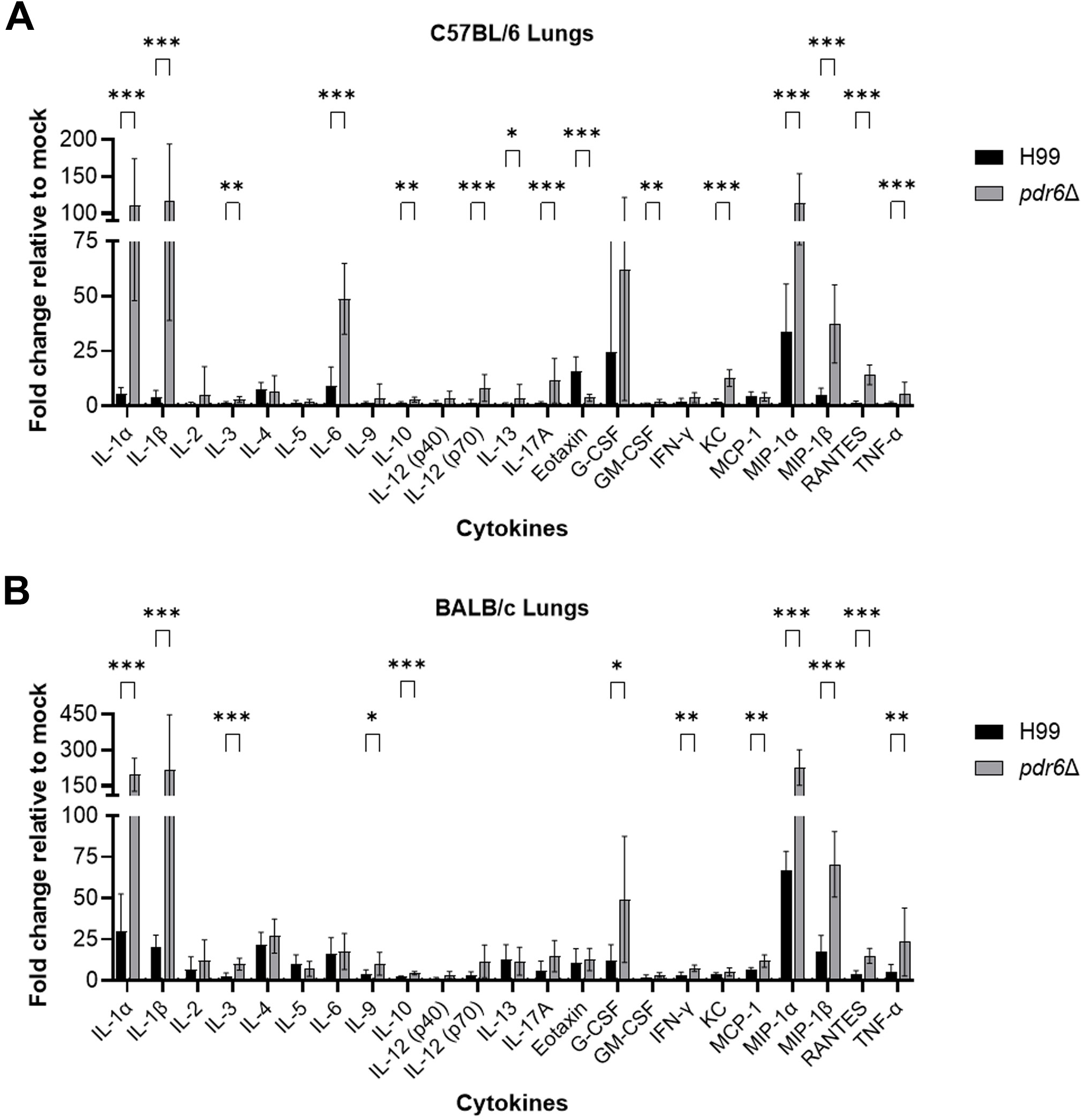
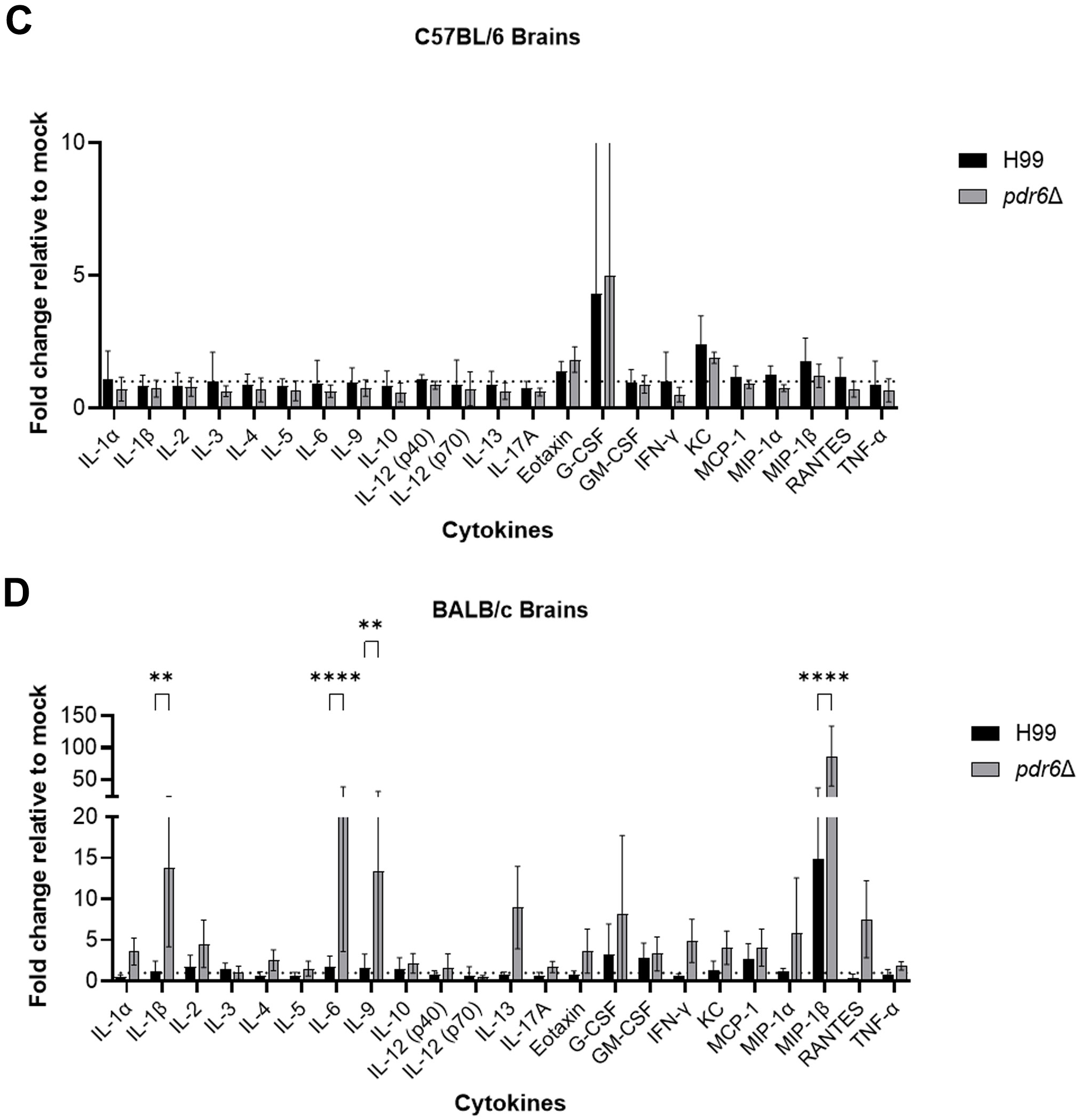

**Figure S5.**
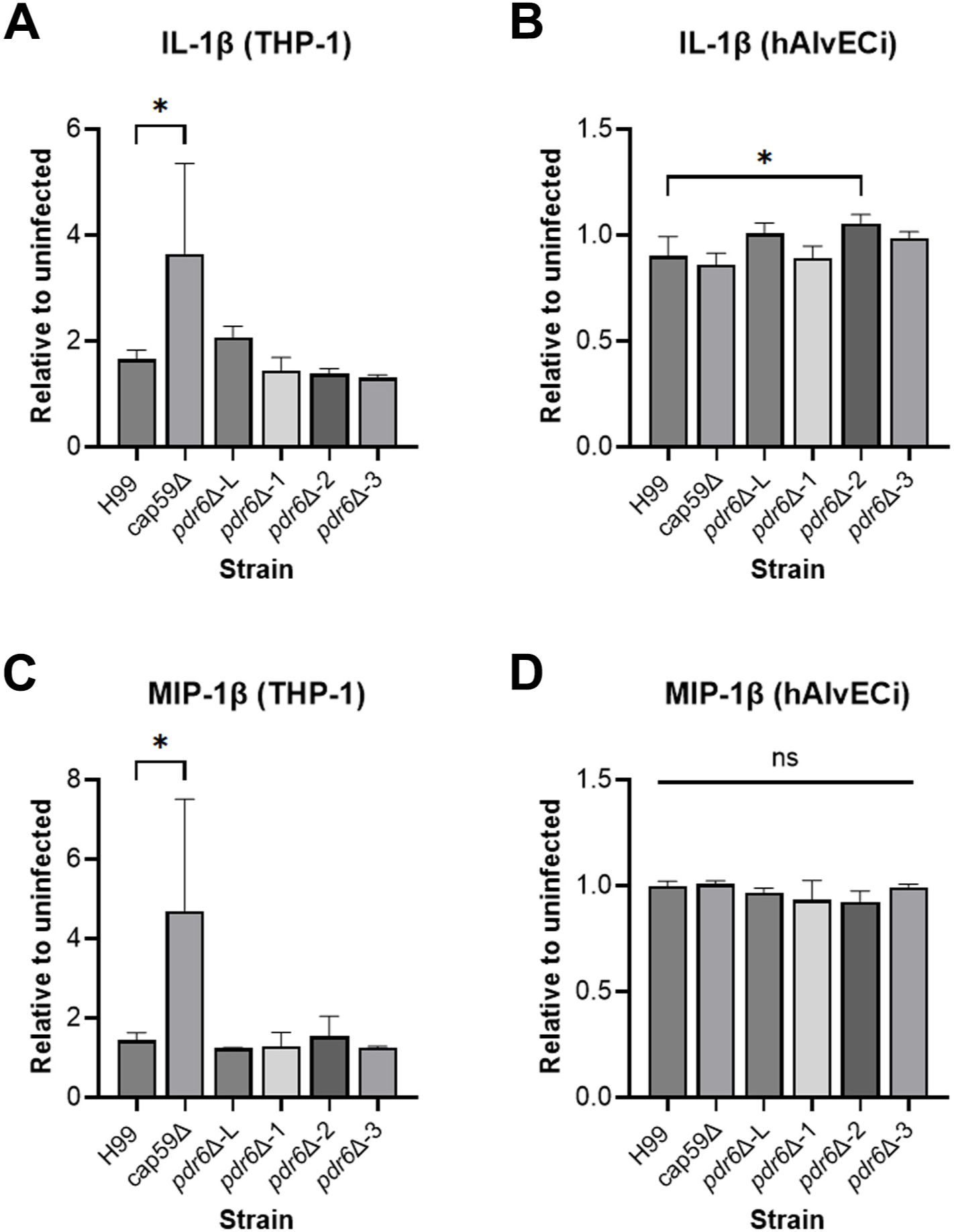

